# Functional imaging of time on task and the involvement of dopaminergic and cholinergic substrates in cognitive effort and reward

**DOI:** 10.1101/2024.12.12.628171

**Authors:** Chiara Orsini, Julia E. Bosch, Karin Labek, Roberto Viviani

**Affiliations:** Institute of Psychology, University of Innsbruck, 6020 Innsbruck, Austria; Department of Psychiatry and Psychotherapy III, University of Ulm, 89075 Ulm, Germany

**Keywords:** “sustained attention”, “time on task”, “cognitive effort”, “reward”, “basal forebrain”, “ventral tegmental area”

## Abstract

Neuroimaging studies have identified the neural substrates associated with sustained cognitive efforts and control and their modulation by rewards. Different lines of evidence implicate the prefrontal cortex (especially the anterior cingulate cortex, ACC), dopaminergic, and cholinergic substrates in this modulation. We studied here the activity of these substrates at increasing time on task (requiring increasing levels of cognitive effort) in trials within blocks with differing reward levels. In the cortex, while peaking in the ACC, activity associated with time on task was extensive, also including activity decrements outside the default mode network, primarily involving motor and somatosensory regions. Information about reward levels was carried in the ventral striatum, consistent with its motivational role, but did not reflect trade-offs with increasing efforts during time on task. Instead, the ventral tegmental area and parts of the basal forebrain (BF) corresponding to the cholinergic Ch4 nuclei increased in activity with time on task and were sensitive to reward levels. This BF activity is consistent with a cholinergic role in driving compensatory efforts modulated by reward levels. These findings identify the BF as a neuroimaging phenotype associated with sustaining task sets and cognitive efforts.

## Introduction

Sustained attention^1–4^ is conceptualized as the cognitive capacity required to remain task-focused over a prolonged time^5–8^. A common observation, also based on experience, is that this requires effort^9^ and that performance may worsen with time^8^. The term “vigilance” is sometimes used in this context, although with less specificity, often also referring to the arousal state^6^, which may degrade with prolonged time on task (ToT). However, Posner and Boies argued that sustained attention is already involved during relatively brief tasks^10^, a view supported by meta-analytic evidence on its recruitment after ten seconds of work^11^. A seconds-lasting ToT may be specifically informative of the sustained attention processes required to compensate momentary lapses of goal-directed attention, while prolonged ToT induces mental fatigue, i.e. vigilance decrements that are associated with a slow increase of error rates^12^. Here, as in these references, we use the notion of sustained attention to refer to effortfully and continuously attending to a short and simple task with little or no working memory load, using a short ToT to avoid the additional burden of mental fatigue and exhaustion arising from several minutes or hours of labor.

The present functional imaging study aims to investigate the neural correlates of sustained attention and the effect of reward levels on these potential correlates by looking at the effects of short ToTs while also considering the possible involvement of dopaminergic, noradrenergic, and cholinergic substrates. This focus addresses issues raised in separate strands of theory and evidence in the literature on sustained attention and cognitive effort and their modulation by reward, possibly referring to interrelated mechanisms. To our knowledge, these separate strands have never been investigated within the same framework in humans. Furthermore, no functional imaging study has investigated the effects of changes in reward levels in the sustained attentional processes observed at short ToT, focusing instead on tasks with high working memory load.

In the present experiment, a simple detection task was modulated by varying levels of reward^13^. A cue, announcing the level of reward obtained by performing the task correctly, was followed by a “foraging patch” of about 15 seconds, where participants had to press the right or the left button each time a stimulus appeared at the right or left side of the screen to collect the announced reward (Figure 1A). Previous characterizations of this task^13^ have shown activity of the ventral striatum depending on the lack of predictability of the valence of the cue, as in well-known prediction error models of dopamine signaling^14,15^ and as in numerous previous neuroimaging studies^16–23^. The foraging patch, in contrast, carries no prediction error but is characterized by different levels of reward accruing from doing the task. Reward levels in the foraging patch were associated with activity in the ventral tegmental area/substantia nigra (VTA/SN), as well as in the nucleus accumbens (NAcc, Figure 1C)^13,24^. In the present study, we investigated short ToT effects in the foraging patch using a database of N=415 individuals that completed this task in the scanner^24^. ToT was modeled as the progression of the trials within the 15s blocks (Figure 1B). The resulting regressor is orthogonal to the task-based regressors used in the previous work and provides new information on the neural substrates modulated by this task.

**Figure 1:**
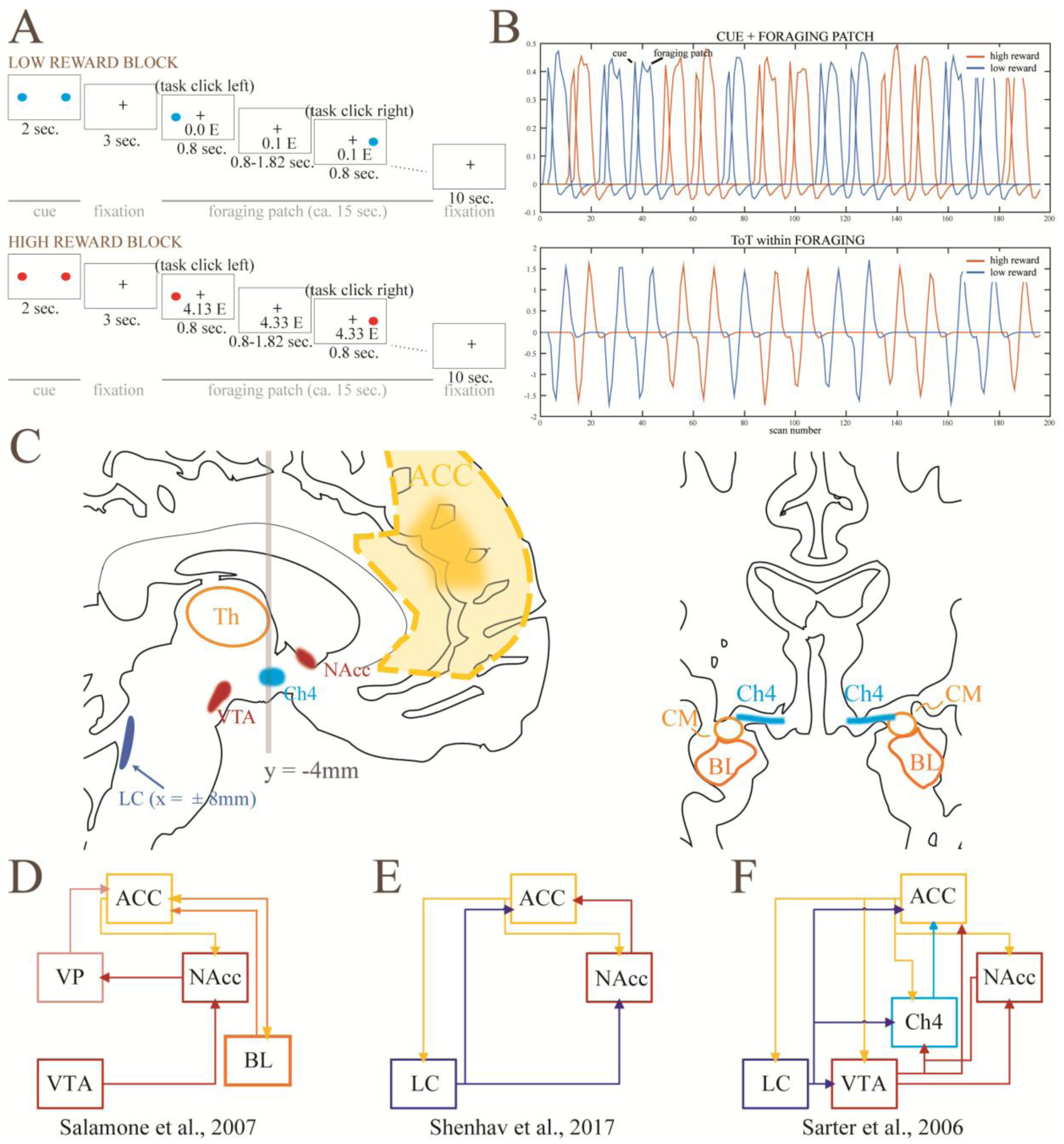
A and B: Schematic representation of the paradigm of the study, consisting of trial blocks with high and low reward levels (“foraging patches”, in blue and red in the figure). A linear trend within the foraging patches indexed time on task (modified from ref.^13^). C: Brain structures involved in models on the interaction between reward and cognitive efforts. D: In the motivational model by Salamone and colleagues, redrawn from ref.^33^, dopaminergic substrates (NAcc and VTA) energize cognition through modulatory activity in the cortex together with the basolateral amygdala (BL), with a preferential distribution to prefrontal areas through the intermediation of the ventral pallidum (VP)^33^. E: In the model of the computation of required effort by Shenhav and colleagues^26^, the ACC signals increased effort requirements to the locus coeruleus (LC) and the NAcc, which modulate cortical activity. F: Cholinergic model of cognitive modulation from the animal literature by Sarter and colleagues, redrawn from ref.^25^. Cholinergic modulation (Ch4) flanks the modulation from LC and dopaminergic substrates^25^. CM: centromedial amygdaloid nucleus.

To interpret these substrates, we draw on theoretical models of effort. Despite its wide use in literature, attentional effort has never been operationalized and investigated as a unique construct^25^. Early empirical investigations of the notion of effort took place within the context of bottleneck or limited resources models of controlled processes more generally^9^. More recently, the notion of effort has been used to model mechanisms that modulate recruitment of cognitive control^26^. Empirical evidence demonstrates that cognitive efforts are avoided in the presence of alternatives, suggesting that cognition internally maintains a representation of its costs^27,28^. This notion is consistent with the observation that rewards attached to the correct execution of tasks affect the levels of effort associated with attentional demands and improve performance (for a review, see ref.^29^), as if rewards were traded off with the costs of cognition for their procurement. This trade-off is apparent also in sustained attention tasks, as perceived effort during continued performance is less when the benefits associated with it are greater^30^. Among all areas activated by executive control, it has been proposed that the anterior cingulate cortex (ACC) is concerned with evaluating the allocation of cognitive effort based on a computation of costs and benefits^26,31^. Many findings in the literature on ACC recruitment are explained by the resulting model, which predicts higher activity in ACC with higher cognitive demands and higher reward levels^26,32^.

By tracking a signal of the reward expected from work, activity in dopaminergic regions during the foraging patch may represent the benefit of allocating cognitive control and the motivational effects of reward^13^ consistently with well known models of the motivational role of dopamine (Figure 1D). In turn, the effort associated with sustained attention may increase during the progression of the foraging patch, representing increasing costs of control and/or a mechanism countering attentional decrements. When looking at the neural substrates that increase with the time on task, we expected to replicate the involvement of the sustained attention cortical network reported in the literature^11^ and the key role of ACC in registering changes in reward levels and cognitive demands in ToT^26^. However, the study also gave us the opportunity to investigate the role of subcortical nuclei mentioned in separate strands of literature on cognitive effort and reward, including animal literature, that have not been systematically considered in this setting.

One question was the change in activity of VTA/SN and NAcc during ToT. Several proposals have been made about the nature of cognition costs^26^. In the case of prolonged but simple tasks, a possible contributor may be “opportunity costs”^34^, i.e., the notion that remaining on the same task for a long time implies forfeiting returns that may be obtained with less effort elsewhere, a computation not unlike the one formalized by foraging models^35^. Previous neuroimaging studies have shown that activity in dopaminergic substrates is decreased by cognitive efforts to obtain rewards^36,37^, although the animal literature is not supportive of this finding^38^. We hypothesized that VTA/SN and NAcc, activated here by levels of reward, would also track the diminishing net value of the rewards during the sustained attention period.

A further issue discussed in the neuroimaging literature on the ACC concerns the recipients of its output, an issue related to the mechanisms through which control is modulated. A working hypothesis is the involvement of the locus coeruleus (LC)^26^ (Figure 1E), due to evidence on its role in modulating attention^39,40^. In this literature, this model also includes the ACC receiving input from the anterior insula to implement increased cognitive engagement^26,32^.

However, in the animal literature simple signal detection tasks have also provided abundant evidence that cholinergic activity from the nucleus basalis of Meynert (NBM) Ch4 region located in the basal forebrain (BF)^41–43^ mediates sustained attentional performance (for reviews, see ref.^44–46^). Cortical cholinergic activity increases in cases of cognitive challenges, including ToT, and is thought to compensate for the possible reduced attentional performance during these challenges and to be key to the instantiation of cognitive effort^25^. Interestingly, the animal literature has also provided evidence of the sensitivity of the cholinergic BF to reward contingencies^47,48^, and that incentives have an immediate supporting effect on cognitive efforts^25^, providing yet another possible avenue for the computations involved in efforts to sustain attention (Figure 1F).

Our study provided two novel insights. The first was that, while the expected cortical activations associated with cognitive effort were replicated in the data, a sample of this size also provided evidence (at voxel-level correction) that most of the cortex was modulated by ToT in one way or another. In our data, the cortical activations reported in the literature looked like the tip of the iceberg and were opposed to isolated decrements of activity with ToT, not located within the default mode network.

The second was that cholinergic BF and VTA were active and modulated by reward in a manner dissociated from NAcc. There were no effects in the brainstem in the location of the LC. This finding suggests that, in a very simple task, cholinergic activity may play an important role in the maintenance of the task set in humans, while the motivational value of reward is tracked by a signal emerging in the ventral striatum.

## Results

### Behavioral data

Behavioral data were reported in a previous study^24^. Briefly, participants could carry out the task with ease (giving the correct response in over 99% of targets). After adjusting for the first trial in block, where participants were significantly less accurate (*z* = -7.9, *p* < 0.001), correct responses diminished with the ToT (*z* = -4.2, *p* < 0.001). The reward amount influenced accuracy as well, with worse performance in the less rewarding condition (*z* = -3.6, *p* < 0.001).

Reaction times (RTs) were shorter in high reward foraging blocks by about 3.2 ms (*t* = 9.1, *p* < 0.001). After adjusting for the first trial, which was slower than the subsequent trials (of about 84.6 ms, *t* = 117.5, *p* < 0.001), RTs showed increases during the foraging block, i.e., a ToT effect of about 0.3 ms per trial (*t* = 4.5, *p* < 0.001). The significance of the ToT effect was robust to variations of the model. For details, see the R Markdown section in the Supplementary Materials.

### Neuroimaging data

In the following, 8 mm smoothing was used to analyze cortical effects and in the Results Tables in the Supplementary Materials. In the text, we report on an analysis at 4 mm smoothing for effects in subcortical nuclei and the amygdala. In both cases, significance levels were corrected at voxel level (peak level) for the whole brain volume.

### Effect of reward during the foraging phase

The main effects of reward levels were described in ref.^13,24^ and were replicated here. Briefly, the effect of reward in the foraging phase showed an involvement of VTA/SN (x, y, z: 7, -13, -10, *t* = 6.66, *p* < 0.001), and of the NAcc (-7, 5, -4, *t* = 9.46; and 6, 5, -4, *t* = 7.84, all *p* < 0.001 voxel-level corrected). Of note, no effect of reward was observed in the Ch4 or in the LC in this contrast.

### Effect of ToT

The effect of ToT was modeled by the linear time during the foraging blocks, when participants executed target detection continuously. This model revealed widespread cortical and subcortical activity increases with increased ToT (Table S1; Figure 2, red-yellow colors). The largest cluster was localized in prefrontal areas (both medial and dorsolateral), and extended to parietal, temporal, occipital, and subcortical regions.

**Figure 2:**
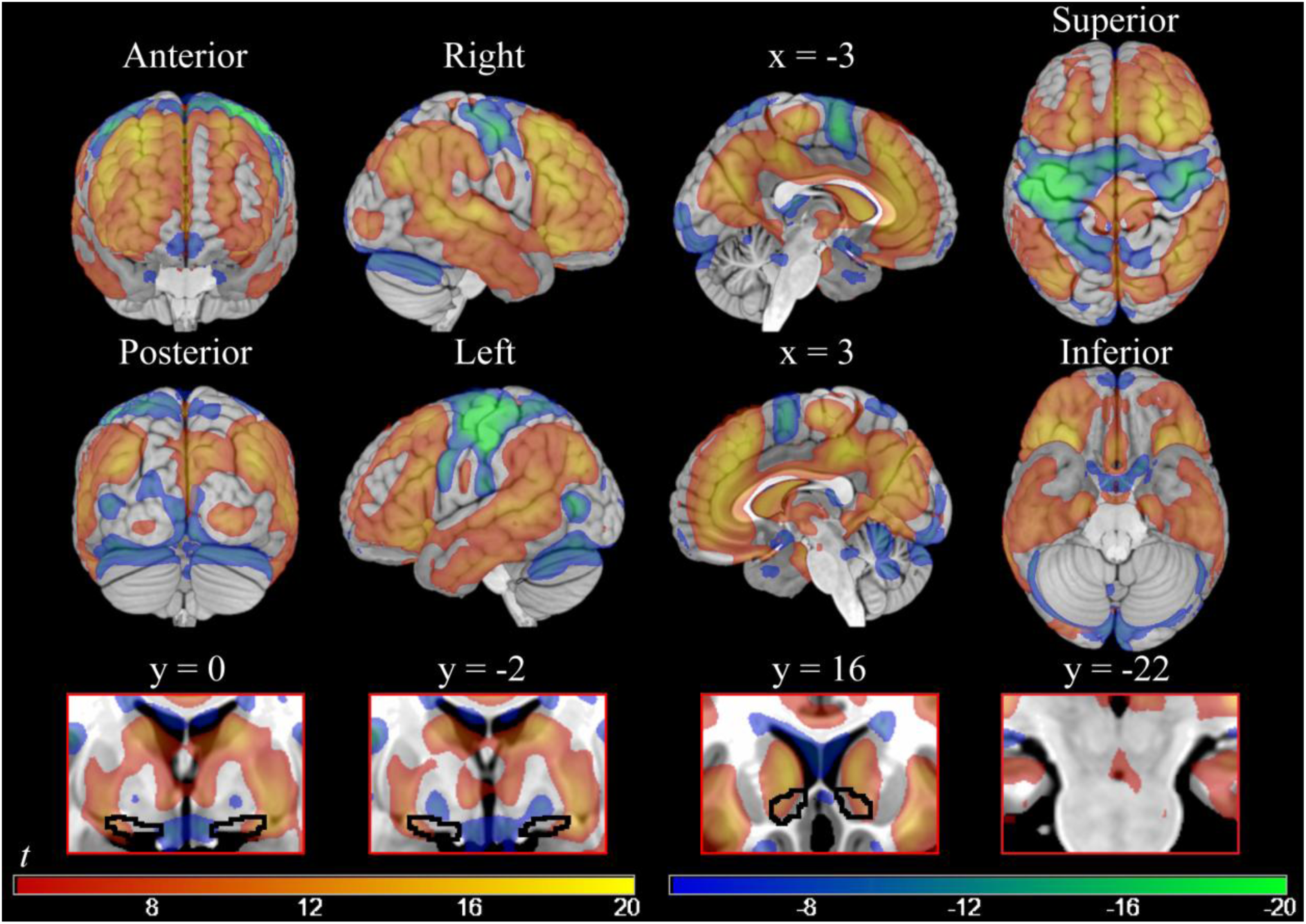
On the top, parametric maps of voxel-level significant cortical activations and deactivations associated with increasing time spent on task (smoothing = 8 mm). On the bottom, subcortical significant activity at the same threshold (smoothing = 4 mm) in Ch4 (y = 0; y = -2), NAcc (y = 16), and VTA/SN (y = - 22). Areas showing an activity increase are the ones in the red-yellow gradient; areas showing an activity decrease are the ones in the blue-green gradient. T-values = ±4.5/±20.

Some cortical substrates showed activity decreases with ToT (Table S2 and Figure 2, blue-green colors). Particularly, the areas showing this pattern bilaterally (even though presenting a stronger deactivation in the left hemisphere) were the premotor/motor and somatosensory cortices, and the left associative visual cortex. In separate analyses (not reported for brevity), we verified that these decreases were present in models without adjustment for “physiological noise” confounders (data available upon request).

In subcortical regions, increases in activity with ToT included the basal ganglia, the thalamus (x, y, z: 6, -10, 6, *t* = 10.73), the central nucleus of the amygdala (centromedial amygdaloid area, 25, -1, -11, *t* = 14.09, and -24, -1, -11, *t* = 11.81), the lateral portions of the cholinergic Ch4 NBM component (24, -1, -10, *t* = 13.20, and -22, -1, -11, *t* = 11.61), the VTA/SN (3, -16, -10, *t* = 8.63), and the NAcc (-12, 15, -4, *t* = 12.48, and 13, 17, -5, *t* = 10.82; all at voxel-level significance *p* < 0.001 with 4 mm smoothing; see the bottom row of Figure 2). Hence, the subcortical areas that we hypothesized as being related to ToT were significantly involved. However, the changes in VTA/SN and NAcc activity over time were in the opposite direction as in our *a priori* hypothesis. Furthermore, activity in the NAcc and Ch4 was embedded in a large activation blob, making evaluation of its specificity more difficult. NAcc, for example, was part of a general activation of the caudate. In subcortical regions, ToT was associated with activity decreases in the pallidum (-15, -4, -8, *t* = -12.67, and 16, -2, -7, *t* = 9.58; voxel-level significance *p* < 0.001 with 4 mm smoothing, see insets in the bottom row of Figure 2 at y = 0 and -2).

### Interaction ToT x reward level

The interaction between ToT and reward levels revealed a mostly right-lateralized pattern of activation (Table S3; Figure 3, red-yellow colors) in areas that were recruited also in the ToT analysis (Figure 3, shaded yellow layer). Hence, in the major cortical areas where activity increased during ToT it did so more in high reward trials, although this effect was more markedly right-lateralized. A notable exception to this lateralization pattern was ACC/medial prefrontal cortex, where the highest effects of the interaction were reached (black asterisk in Figure 3 at *x* = 3).

**Figure 3:**
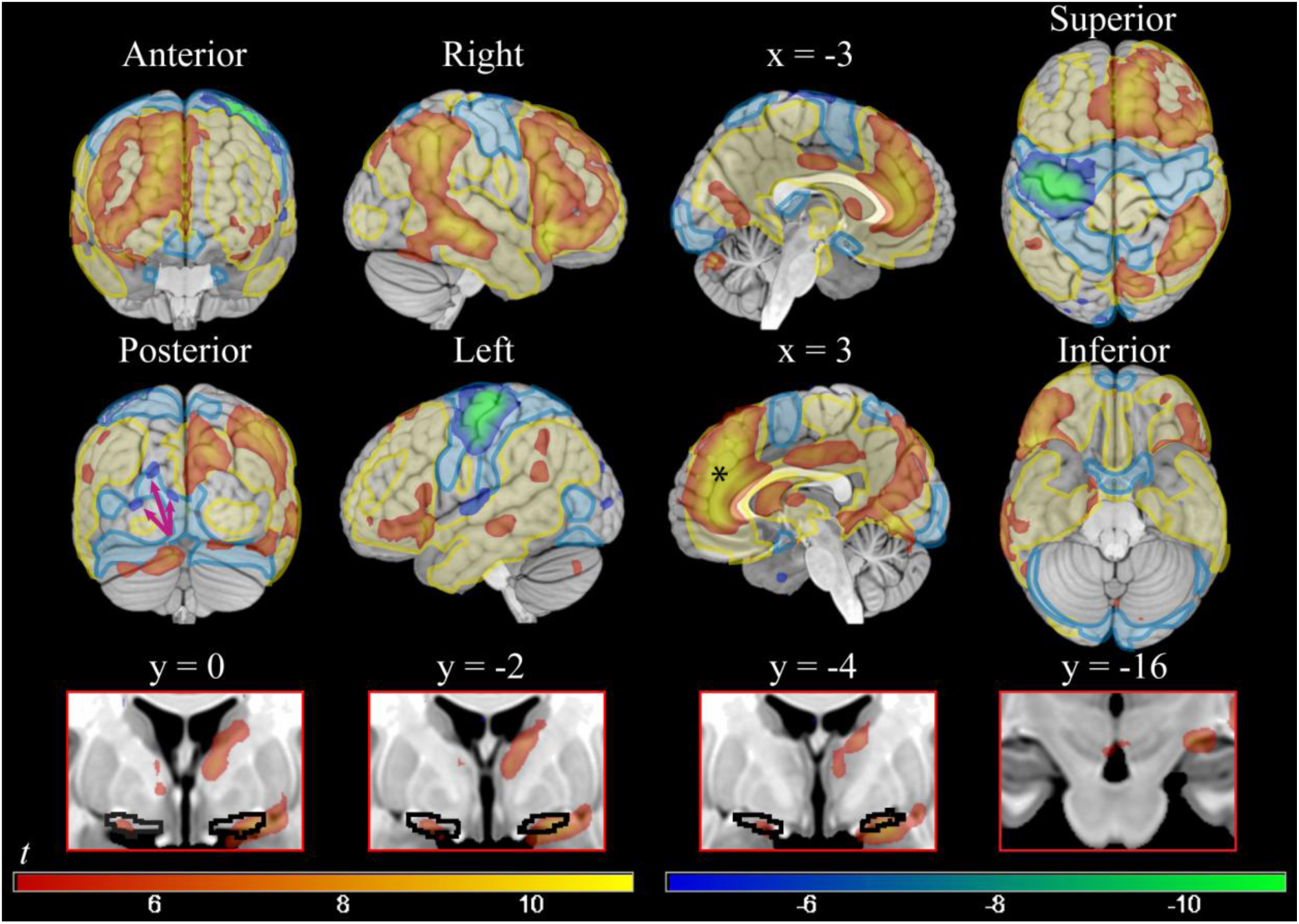
On the top, cortical voxel-level significant parametric maps of the positive and negative interaction between time on task and levels of reward (high reward vs low reward) (smoothing = 8 mm). On the bottom, subcortical significant activity at the same threshold (smoothing = 4 mm) in Ch4 (y = 0; y = -2; y = -4), and VTA/SN (y = -16). Areas shown in red-yellow presented greater activity during time on task in high than in low reward conditions; interaction in the other direction is shown in blue-green. T-values = ±4.5/±11. On the top, in shaded yellow and blue, positive and negative activations in the time on task of Figure 2, showing that areas modulated by reward in the interaction were mostly a lateralized subset of the time on task effects. The black asterisk at the sagittal slice at *x* = 3 shows the peak effect of the interaction at *y* = 41, *z* = 23.

In the opposite direction (Table S4; Figure 3, blue-green colors), more marked decreases in ToT during high reward trials involved an almost totally left-lateralized set of fronto-parietal areas, which overlapped with the fronto-parietal deactivations in the ToT contrast (Figure 3, shaded blue layer). Also, the superior occipital cortex showed significant decreases (arrows in the posterior view in Figure 3).

While not as extensive as in the ToT contrast, also the interaction with reward involved large cortical regions with internal peaks of activity. To visualize these peaks, in Figure 4A we show the interaction at a very high threshold level. In Figure 4B and 4C we show the areas of connectivity and co-activation of the seed of the ACC peak, taken from the Neurosynth database (www.neurosynth.org), showing that in both our data and in the database, ACC was connected with the other peaks, including that of the anterior insula, and presented with co-activations in the middle temporal gyrus and inferior parietal lobe. In Figure 4D, the decoding analysis from the Neurosynth database revealed that these areas were active in many studies in the literature involved with cognition, where the association referred to cognitive conflict, although the strongest overlap involved social cognition tasks.

**Figure 4.**
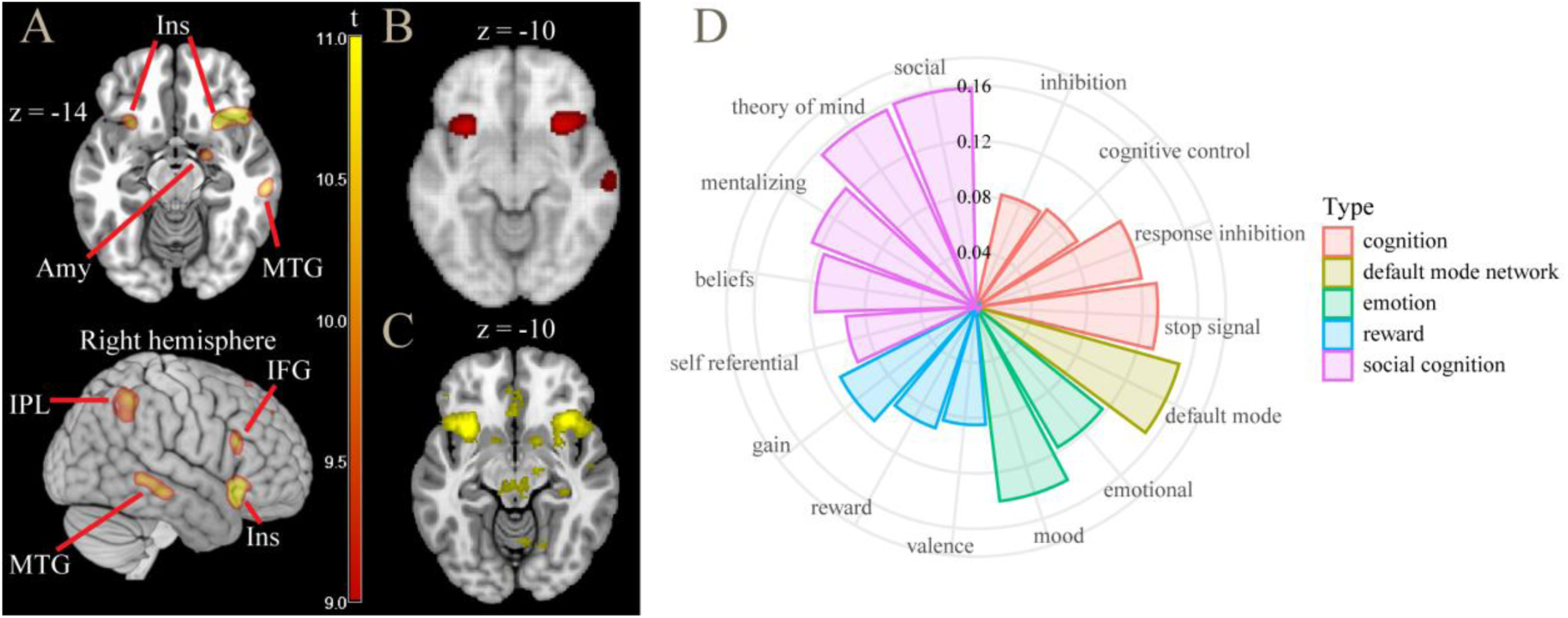
A: contrast time on task × high vs low reward: areas active at high threshold levels (*t* > 9.0). B: connectivity map of ACC seed, showing high co-activation in anterior insula. C: meta-analytic co-activation map (*z* > 4.0), showing co-activation in anterior insula and central nucleus of the amygdala. D: decoding analysis of this contrast (from www.neurosynth.org). ACC = anterior cingulate cortex; Amy = amygdala; IFG = inferior frontal gyrus; Ins = insula; IPL = inferior parietal lobule; MTG = middle temporal gyrus.

Subcortically, similarly to the result in the ToT contrast, activations were detected in the NBM/Ch4 (peaking bilaterally in x, y, z: 21, -4, -11, *t* = 9.19, and -19, -1, -13, *t* = 6.40, all *p* < 0.001, here and in the following voxel-level corrected at smoothing 4mm), also including the centromedial amygdaloid nucleus (21, -4, -13, *t* = 9.24, *p* < 0.001, and -19, -2, -14, *t* = 5.29, *p* = 0.015). Differently from the activations found in the ToT contrast, these regions included both lateral and medial components of the Ch4 and presented a more specific localization. Also VTA/SN was involved, showing a small but significant effect (3, -14, -13, *t* = 5.15, *p* = 0.025), as well as the thalamus (9, -1, 6, *t* = 6.96, *p* < 0.001). There was no significant interaction between ToT and reward levels in the NAcc and in the LC (4, -36, -19, *t* = 2.00, *p* > 0.05, all voxel-level corrected at smoothing 4mm). Ribbon plots showing the course of relative activation in the foraging patch for these regions are in Figure 5.

**Figure 5.**
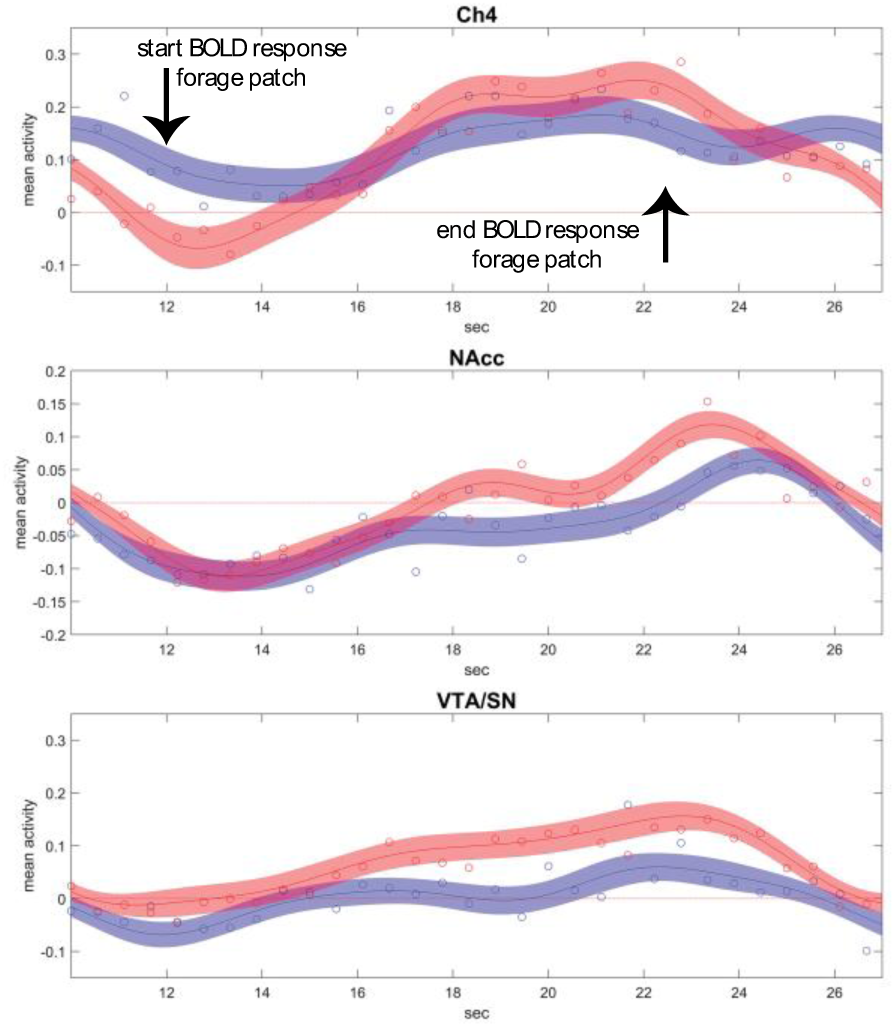
Ribbon plots showing relative activation courses in Ch4, NAcc, and VTA/SN (from top to bottom, fitted Fourier series and 5% confidence intervals) during the high (in red) and low (in blue) reward foraging patches. The circles are median relative activity at the sampled time points. In Ch4, activity during low reward blocks shows only a small increment, in contrast to the high reward blocks. NAcc shows consistent increases, larger in the high reward blocks, but not enough to survive correction in the statistical analysis. The VTA/SN activity is characterized by higher sustained activity in the high reward blocks, but also stronger increase during the progression of the block. Data adjusted for the constant term, movement covariates, and physiological noise confounders.

In summary, the interaction between ToT and reward revealed that at high reward levels both increases and decreases of cortical activity associated with the ToT were essentially a more lateralized subset of those of the ToT effect. In the BF, the involvement of areas consistent with recruitment of Ch4 was more specific than in the main effect. In the midbrain, the VTA exhibited a small but significant sensitivity to higher levels of reward.

## Discussion

### Behavioral effects of ToT and reward

ToT and reward levels had very small but significant effects on participant’s performance. While ToT negatively impaired accuracy and RTs, reward positively modulated these measures. This finds confirmation in the literature on the behavioral effects of ToT and reward^49–52^. Furthermore, the very small size of the effect of reward levels suggests that the need to reach a threshold to obtain the reward forced participants to apply constant effort during the task irrespective of reward levels.

### Cortical effects of ToT and its interaction with reward levels

The recruitment of predominantly right-lateralized prefrontal and parietal areas of the present study has been widely associated with sustained attention and cognitive control in the literature^5,12,53,54^; for a systematic review and meta-analysis, see ref.^11^. The functional involvement of thalamus, also present here, is frequently reported in sustained attention literature^5,11,55^.

Similitudes aside, there were also differences between our results and those of previous studies, as summarized in ref.^11^. The areas recruited by ToT (Figure 2) were embedded in a much more extensive modulation of cortical activity. Instead of presenting as discrete specific areas, the effects of ToT may be better described as gradients of activation peaking in anterior and posterior cingulate cortex, the anterior insula, and the frontal and superior parietal foci. Because of the empirical evidence of the effect of reward on cognitive effort^29^, it is the interaction between ToT and reward levels that may throw light on the substrates specifically involved in modulation of control. We found that the areas most involved with ToT were also modulated by reward levels (Figure 3), supporting their role as the neural correlates of the energization of attentional processes by reward^29,51,52,56–58^. Among these areas, the ACC and the right anterior insula stood out by presenting the largest effects. Comparison with rest connectivity data revealed these areas to form a highly connected network, which may constitute the core of the neural mechanism for the modulation of control in association with reward levels.

This finding is consistent with the notion that ACC computes signals tracking the cognitive control demands required to maximize the difference between expected benefits and intrinsic costs of cognitive efforts^26,32,59^. The same model attributes to the anterior insula the role of providing input to the ACC about salient events in the environment requiring control adjustment, and the involvement of the lateral prefrontal cortex, also active here, in the implementation of the control process specified by the ACC^59^.

A novel finding was that there were also cortical areas where activity decreased with ToT. These were the motor, premotor, and sensory cortex (extending ventrally into the parietal operculum) and left visual association areas. These decrements were more marked in high reward trials, arguing for their functional nature. In the neuroimaging literature, decreases in cortical activity have been interpreted as neural correlates of priming effects^60–62^. Priming in neuroimaging studies has been shown to ensue from diverse mechanisms. Perceptual or neural priming is typically localized in associative visual or auditory areas^63,64^. This may explain the decrements in the left visual association areas observed here. Activity decrements in frontal areas have been attributed to conceptual priming^65,66^, but the prefrontal areas described in these studies did not show decrements in our data.

In the present study, the largest activity decreases, located in the sensorimotor cortex, may not be consistent with perceptual priming, given the visual nature of the stimuli. However, decrements in the sensorimotor cortex during the progression of the trials may be observed for example in decision making (where participants decide between options presented on the right and the left with a button click, as was the case here) as part of a larger pattern of activity decreases that include visual associative cortex^67^. Decreases in activity with ToT during passive exposure to visual stimuli similarly involve visual associative cortex, while sparing the sensorimotor areas^68^. In the present study, the associative visual cortex was also involved, although to a lesser extent. We speculate that the very simple task of the present study may explain the regionally limited involvement of activity decreases in visual associated cortex, relative to what observed in a decision-making task with more complex stimuli, and the more marked involvement of sensorimotor areas.

### Recruitment of VTA/SN and NAcc

We hypothesized that VTA/SN activity may decrease during the foraging patch and encode the diminished returns of the task at the net of the “opportunity cost” of rewards from alternative courses of action, or the increased cost of maintaining the task set. Contrary to our hypothesis, activity in VTA did not decrease during the foraging patch, showing instead a modest but significant increase.

There are several possible reasons for the discrepancies between our result and previous observations of decreased activity in dopaminergic substrates representing outcomes discounted by effort^36,37^. First, the results in the paper by Botvinick and colleagues^36^ were obtained by looking at differences between obtained and expected outcomes (prediction errors), a signal that in our paradigm arises at the cue, not during the foraging patch. Indeed, the paradigm was designed to disentangle between motivation triggered by reward-predicting cues and the motivation to work to obtain expected rewards^13^. Second, the areas identified by Kurniawan and colleagues^37^ were in the ventral putamen and pallidum and were more posterior than the NAcc. Our data did contain ToT activity decreases in the pallidum. However, it appeared to be located more posteriorly than the activity reported by these authors, making its interpretation uncertain.

The modulation of dopaminergic substrates we observed in the interaction between reward and ToT differed from those induced by levels of reward in the task, because it was limited to the VTA/SN and did not involve NAcc. In the ToT contrast, NAcc activity increased. However, this was part of a general activity of the whole caudate. Our findings confirm the involvement of NAcc in the increased energization of responses in the high reward trials^69^, but not in the effects of reward on ToT activity. These findings are consistent with those from the animal literature, which suggest that the modulation of the dopaminergic system by expected effort requirements is weak and inconstant, in contrast to the modulation associated with prospective rewards^38^. Unlike NAcc, VTA/SN (together with NBM, see below) was sensitive to the progression of the foraging blocks at different reward levels, during which tracking the rewards accrued by the foraging activity represents a computation of possible relevance to remaining on task.

Recent data on laboratory animals have uncovered several populations of neurons in the VTA^70^, whose diverse functions are only beginning to be understood^71,72^ but include tracking the value of expected reward as is required to compute changes in reward expectancies (prediction error^73^). Unlike dopamine neurons, a GABA subpopulation has been shown to fire irrespective of cues signaling changes in reward expectancies, tracking the value of reward during the consummatory phase instead, and to project to pallidal neurons coding reward-dependent motor activity^74^. By observing the effect of stimulating this VTA-pallidal circuit, these studies show it to be instrumental in maintaining motivation to work for rewards. Furthermore, this activity is dissociable from the dopaminergic drive in NAcc^75^. NAcc shell activity is modulated by VTA GABA neurons, resulting in increased place preference and operant responding^76^. A recent study has shown that VTA activity in these pathways correlated with the response latency or other indicators of intensity of consummatory behavior^77^. While our understanding of VTA is still too incomplete to draw conclusions on the interpretation of our findings, we believe that there are considerable points of contacts between the characteristics of the VTA-pallidal GABAergic circuits of the animal literature and the VTA activity uncovered here and in a previous neuroimaging study^13^.

### Recruitment of LC

Contrary to what hypothesized, there was no activity in the region of the LC in any contrast in our data. Although this null result may be due to a limited spatial sensitivity or a partial volume effect, LC was the only small nucleus that did not show an involvement among those reported in the literature. Despite the isotropic voxel size utilized at data collection (3 mm), the large sample size (over 400 participants) should provide sufficient power to detect significant effects. The heterogeneity of the characterizations and paradigms investigating sustained attention or ToT likely accounts for this discrepancy. Indeed, it has been observed that LC modulates cognition through arousal^78^, whose effects could be better captured in the case of longer paradigms.

### Recruitment of putative cholinergic substrates in the BF

The involvement of cholinergic BF in cognition is supported by the abundant evidence in sustained attention paradigms in animal studies^79–84^. In this literature, NBM cholinergic activity increases in the presence of cognitive challenges, i.e. factors that hinder the regular execution of the task, of which ToT is one example^25^. About 90% of the NBM consists of cholinergic neurons known as Ch4^41,42,85^, which constitute the main source of cholinergic projections to the cortex^42^.

In contrast to these animal studies, the recruitment of cholinergic components of BF has been rarely reported in sustained attention human studies (for an exception, see ref.^86^). It should be emphasized that in our findings Ch4 recruitment became spatially specific only in the interaction between ToT and reward levels. The effects of ToT alone in Ch4, while not incompatible with its recruitment, were not conclusive due to the enmeshment of this region in a stronger regional pattern of activation.

Our findings are consistent with those of animal studies reporting the involvement of NBM when processing reward-related information, as in reward anticipation^48,87^, reward/punishment delivery^47,48^, and cue-reward learning^88^. In a sustained attention paradigm in mice, Tashakori-Sabzevar and Ward demonstrated the BF’s involvement in converting the motivational importance of probability signals of reward into enhanced attentional performance^89^. As argued by Sarter and colleagues^25^, cholinergic BF may play a role in the modulation of increases in attentional effort, in particular in the presence of cognitive challenges and when motivated by a rewarding outcome. In this literature, cholinergic activity is often considered as supporting cognition in the face of challenging conditions, a compensatory mechanism distinct from motivation. Our data support the distinction between mechanisms involved in motivation and those involved in sustaining effort related to rewards, as NAcc was not recruited in the interaction between ToT and reward levels, in contrast to VTA/SN and NBM.

Our finding of the selective involvement of NBM/Ch4 in the interaction with reward levels may open the way to the use of this neuroimaging phenotype in future studies of individual differences^90^. For example, subsequent work has shown that during the sustained attention task, NBM/Ch4 activity increases with ToT are positively associated with individual differences in self-regulation^91^. This finding further supports the notion that cholinergic BF function is critically involved in sustaining attentional effort, and may associate with individual characteristics.

## Conclusion

The cortical effects of ToT confirmed the results of previous studies of cognitive effort and sustained attention and showed that varying levels of reward may modulate these effects consistently with the facilitatory effects of reward on cognitive effort. However, our effects also included areas where activity decreased with ToT, such as the premotor/motor and somatosensory cortex, and involved most of the cortical mantle. We found no evidence of involvement of the LC in any of the contrasts we examined.

We found no evidence in support of our initial hypothesis that dopaminergic substrates may signal diminishing levels of perceived returns, after discounting increasing effort while remaining on task. Instead, ToT was associated with increasing NBM/Ch4 activity, which was consistent with the abundant evidence for the involvement of this nucleus in the animal literature. This activity was best identified in the modulation of ToT by reward, together with activity in VTA/SN.

This finding suggests a partial dissociation of dopaminergic and cholinergic activity in the modulation of cognitive control. NAcc, which is widely associated with the motivational effects of reward and was here more active in high reward trials, was not involved in the modulation of the ToT by reward levels. In contrast, this modulation was associated with NBM/Ch4 and VTA/SN activity, which may identify a distinct phenotype involved in the maintenance of task sets and cognitive effort in interaction with a cortical network centered on ACC and the anterior insula.

## Materials and methods

### Participants

The data used in the present study were collected as part of a genetic imaging research project^24^. Written informed consent was obtained from all participants. 441 healthy participants were screened for psychiatric disorders, and they were excluded if they presented current substance (alcohol or drug) addiction, anorexia, present affective disorders, pregnancy or breastfeeding, severe acute or chronic illness, assumption of psychoactive or long-term medications, presence of metal implants, large tattoos, or tattoos near the head. Further exclusion criteria in later stages of the study (imaging scan or data analysis) were the presence of clinical findings, artifacts or excessive movements, and equipment or task administration/completion failures. The final sample consisted of 415 subjects (234 females, age range 18-45, mean age 23.43 ± 3.80 years). Data of 58 participants were collected at the German Center for Neurodegenerative Diseases (DZNE) in Bonn, the one of 357 were collected at the University of Ulm.

The study (acronym: BrainCYP) was registered in the German Clinical Trials Register (DRKS-ID: 00011722), followed the guidelines of the Declaration of Helsinki, and was approved by the Ethical Committee of the University of Bonn (No. 33/15) and the Ethical Committee of the University of Ulm (No. 01/15).

### Experimental paradigm

The paradigm used here (Figure 1A) was introduced in previous studies^13,24^. While inside the MRI scanner, participants executed a sustained attention paradigm consisting of 16 blocks, which were characterized by two possible rewarding levels: high and low. Each block consisted of the 2s presentation of a cue, providing information on the reward level, a 3s fixation cross, and a “foraging patch” (12/13 trials) lasting approximately 15s. During the foraging patch, participants had 0.8s to press a button congruently with the location of a dot appearing on the left or right side of the fixation cross (i.e., the left button in the case of a left-presented dot; the right button in the case of a right-presented dot). The interstimulus interval (ISI) varied according to a Poisson schedule with an average temporal interval of 1.23s. Between the end of one block and the beginning of another, a fixation cross was presented for 10s. Participants collected virtual coins when responding correctly (20 cents per correct response in the high reward blocks, 1 cent in the low reward blocks). The final monetary reward was obtained only if participants had won a total minimum amount of 20 euros. Participants were informed that they would need to provide correct and rapid answers during both the high and low reward trials to reach that threshold. This threshold could be reached only within the final trials of the last block, forcing participants to work hard at collecting virtual coins during the whole task irrespective of reward levels. The task was calibrated in pilot studies to ensure participants maintained rapid responses to maximize reward collection, thus avoiding recruitment of strategic processes during the trials, and ensuring small response time variations. Difficulty was intentionally kept low, such as to avoid possible confounding after-effects of misses and errors. The whole duration of the task was equal to 8 minutes. Participants had the opportunity to practice the task prior to being positioned in the scanner.

### fMRI data acquisition

Neuroimaging data were collected at the German Center for Neurodegenerative Diseases (DZNE) in Bonn and at the Department of Psychiatry and Psychotherapy of the University of Ulm, using respectively a 3 T Siemens Skyra scanner, and a 3 T Siemens Prisma scanner. 64-channels head coils were used, and a T2*-weighted echo-planar imaging sequence was applied (TR/TE of 2460/30 ms; flip angle of 82°; FOV measuring 24 cm; matrix size of 64 × 64 pixels with dimension of 3 × 3 mm in 39 transversal slices, each 2.5 mm thick, and acquired in ascending order with a gap between slices set at 0.5 mm for an isotropic voxel size of 3 mm). Since regions with high susceptibility or areas presenting high iron levels have a shorter T2*, a compensation was performed by progressively reducing the TE by 8 ms from slice 24 to slice 14. This resulted in a TE of 22 ms for the initial 14 slices acquired in the bottom of the volume (see the procedure in ref.^92^). T1-weighted structural images were collected and assessed individually to detect eventual abnormalities.

### Masks of subcortical areas

The cholinergic Ch4 NBM component was identified with the JuBrain Anatomy toolbox^93–96^. The JuBrain Anatomy toolbox was used also for the identification of amygdalar structures^93–95,97,98^. VTA/SN were identified using their probabilistic maps^99,100^. The NAcc was identified using the Harvard Oxford Atlas^101–104^ distributed by FMRIB Software Library https://fsl.fmrib.ox.ac.uk/. The LC was identified using a two standard-deviations-mask^105^.

### Data analysis

Neuroimaging analyses were carried out using the freely available software SPM12 (Wellcome Trust Centre for Neuroimaging, http://www.fil.ion.ucl.ac.uk/spm/) running on MATLAB (The MathWorks, https://www.mathworks.com). After realignment, data were registered to standard Montreal Neurological Institute (MNI) space. In the segmentation step, the spatial priors provided by this software were supplemented with a prior comprising the pallidum, red nucleus, and substantia nigra (as in ref.^13^). After registration to MNI space and resampling to a voxel size of 1.5 mm, two separate datasets were obtained by smoothing the data with Gaussian kernels of 8 and 4 mm FWHM. In the analyses, the 8 mm smoothing dataset was used for the cortex (reported in the Supplementary Materials’ Results Tables section) and the 4 mm for subcortical nuclei and the amygdala (reported in the main text).

At the first level, the task was modeled by four groups of zero-duration onset series, two for the cue events (low and high reward) and two for the trials (the presentations of the individual targets) in the foraging patches (low and high reward). The onsets were then convolved with the canonical hemodynamic response function. ToT was modeled as a parametric modulator given by the number of the trial within the foraging patch, after centering (Figure 1B). As a result, estimates of the foraging effects refer to activity in the middle of the foraging patch, and the parametric modulator coefficient measures the increase of activity during the patch. Centering the ToT predictor also ensured that all model predictors were reciprocally orthogonal, splitting the variance of the signal into uncorrelated components. Data were high-pass-filtered using SPM default settings (128s). Confounding covariates were a linear time trend, the mean signals of white matter, ventricles, and cranial bone^106^ as separate regressors, and the six head movement terms estimated at the realignment step and their second derivatives (12 movement parameters in all).

Contrasts of interest in previous analyses of this paradigm assessed the effect of reward levels at the presentation of the cue and during the foraging patch (high vs. low reward). Here, the contrasts of interest concerned the parametric modulators: the increase in activity during the foraging patch and the differences in these increases between the conditions of high and low reward levels. Contrast images were taken to the second level and tested by permutation (8000 resamples, cluster-defining threshold *p* = 0.001, uncorrected). Contrasts from the 8 mm smoothing dataset were used for the primary analysis of cortical effects in the main text Figures and in the Supplementary Materials’ Results Tables section. Contrasts from the 4 mm smoothing dataset were reported in the main text and were used to assess effects in the LC, thalamus, NBM/Ch4, amygdalae, VTA/SN, and NAcc, identified by masks as described in the section “Mask of subcortical areas”. As with the fMRI data, these masks were also resampled to an isotropic voxel size of 1.5 mm, ensuring a common spatial resolution before their application in the permutation test. The freely available software MRICroN (Chris Rorden, https://people.cas.sc.edu/rorden/mricron) was used to visualize parametric maps.

To exclude a threshold-proximity/reaching effect, which might occur during the last block, an additional set of analyses was computed. For this scope, previous to denoising, the final block was introduced in the model as confounding covariate. This analysis reproduced, although with a lower power, the effects observed in the model without the last block correction and excluded threshold-related effects (data available upon request).

Ribbon plots were drawn by fitting a Fourier series (9 basis functions) to the trial-averaged data in the foraging patch after adjusting for movement covariates, physiological noise, and the constant term, using the fda package (https://www.psych.mcgill.ca/misc/fda/software.html, see reference^107^). Confidence intervals were obtained by fitting the series to the 95% confidence intervals of these averages. Points in the plots are median values of the data used for the fit. The decoding analysis was conducted with the “Neurosynth Image Decoder” function in the Neurosynth database (www.neurosynth.org). This function loads the whole-brain parametric map of t values arising from a contrast of interest and compares the overall spatial distribution of the statistics with that of the other parametric maps of the database. The maps contain keywords referring to the original contrast of interest, allowing a meta-analytic automatic comparison of the activity pattern with the patterns of the images in the database.

Analyses of RTs, accuracy, and plots were generated using the freely available software R version 4.4.0 (R: The R Project for Statistical Computing, r-project.org) using the packages dplyr^108^, tidyr^109^, lme4^110^, and ggplot2^111^. After excluding missed trials, RTs were filtered using a threshold between 0.25s and 0.8s. The model was fitted with the function lmer and included trial number within the block (time on task), first trial indicator, reward level, block number, ISI, side switch of target relative to previous trial, acquisition site, and an individual group variable modeled as a random effect (see the Supplementary Materials’ R Markdown section for robustness analyses R code). Accuracy analysis was carried out with a logistic regression (function glmer) replicating, in terms of predictors, the structure of reaction times model. Here, correct hits were coded as success, and incorrect responses and misses were coded as failures in the logistic regression.

## Acknowledgements and Competing Interest Statement

A preliminary version of this work was presented at the CogBases workshop (Paris, Institut Pasteur, 10-11 October 2023). This work was supported in whole by an ERA-PERMED grant (project ArtiPro) of the FWF Austrian Science Fund (grant number I 5903) [Grant-DOI: 10.55776/I5903] to Roberto Viviani. The authors declare no competing interests. For open access purposes, the author has applied a CC BY public copyright license to any manuscript version arising from this submission.

## Author contributions statement (CRediT)

**Chiara Orsini**: Data curation, Formal analysis, Project administration, Validation, Visualization, Writing – original draft, Writing – review & editing. **Julia E. Bosch**: Investigation, Project administration, Validation. **Karin Labek**: Data curation, Project administration, Validation. **Roberto Viviani**: Conceptualization, Data curation, Funding acquisition, Investigation, Methodology, Project administration, Resources, Software, Supervision, Validation, Visualization, Writing – original draft, Writing – review & editing.

## Data Availability

The datasets used and/or analyzed during the current study are available from the corresponding author (R.V.) on reasonable request and after verifying that the proposed use is consistent with the research purposes participants agreed to in the written informed consent.

## References

1 Davies, D. R. & Parasuraman, R. The Psychology of Vigilance. (London: Academic Press, 1982).

2 Mackworth, N. H. The Breakdown of Vigilance during Prolonged Visual Search. Quarterly Journal of Experimental Psychology 1, 6–21 (1948). 10.1080/17470214808416738

3 Mackworth, N. H. *Researches on the measurement of human performance*. (Med. Res. Council, Special Rep. Ser. No. 268.). (His Majesty’s Stationery Office, 1950).

4 Parasuraman, R. & Davies, D. R. Varieties of attention. (Academic Press, 1984).

5 Coull, J. T. Neural correlates of attention and arousal: insights from electrophysiology, functional neuroimaging and psychopharmacology. Progress in Neurobiology 55, 343–361 (1998). 10.1016/S0301-0082(98)00011-2

6 Oken, B. S., Salinsky, M. C. & Elsas, S. M. Vigilance, alertness, or sustained attention: physiological basis and measurement. Clinical Neurophysiology 117, 1885–1901 (2006). 10.1016/j.clinph.2006.01.017

7. Timmers, D.Chapter Six - Treating Attention Deficits and Impulse Control in Clinical Neurotherapy (eds David S. Cantor & James R. Evans) 139–169 (Academic Press, 2014).

8 Warm, J. S., Parasuraman, R. & Matthews, G. Vigilance Requires Hard Mental Work and Is Stressful. Human Factors 50, 433–441 (2008). 10.1518/001872008x312152

9. Kahneman, D. Attention and effort. Vol. 1063 (Englewood Cliffs, NJ: Prentice-Hall, 1973).

10 Posner, M. I. & Boies, S. J. Components of attention. Psychological Review 78, 391–408 (1971). 10.1037/h0031333

11 Langner, R. & Eickhoff, S. B. Sustaining attention to simple tasks: A meta-analytic review of the neural mechanisms of vigilant attention. Psychological Bulletin 139, 870–900 (2013). 10.1037/a0030694

12. Robertson, I. H. & O’Connell, R.Vigilant attention in Attention and Time (eds Anna C. Nobre & Jennifer T. Coull) Ch. 6, 79-88 (Oxford University Press, 2010).

13. Viviani, R., et al. Signals of anticipation of reward and of mean reward rates in the human brain. Scientific Reports 10 (2020). 10.1038/s41598-020-61257-y

14 Montague, P., Dayan, P. & Sejnowski, T. A framework for mesencephalic dopamine systems based on predictive Hebbian learning. The Journal of Neuroscience 16, 1936–1947 (1996). 10.1523/jneurosci.16-05-01936.1996

15 Schultz, W. Predictive Reward Signal of Dopamine Neurons. Journal of Neurophysiology 80, 1–27 (1998). 10.1152/jn.1998.80.1.1

16 Abler, B., Walter, H., Erk, S., Kammerer, H. & Spitzer, M. Prediction error as a linear function of reward probability is coded in human nucleus accumbens. NeuroImage 31, 790–795 (2006). 10.1016/j.neuroimage.2006.01.001

17 Berns, G. S., McClure, S. M., Pagnoni, G. & Montague, P. R. Predictability Modulates Human Brain Response to Reward. The Journal of Neuroscience 21, 2793–2798 (2001). 10.1523/jneurosci.21-08-02793.2001

18 Li, J., McClure, S. M., King-Casas, B. & Read Montague, P. Policy Adjustment in a Dynamic Economic Game. PLOS ONE 1, e103 (2006). 10.1371/journal.pone.0000103

19 McClure, S. M., Berns, G. S. & Montague, P. R. Temporal Prediction Errors in a Passive Learning Task Activate Human Striatum. Neuron 38, 339–346 (2003). 10.1016/S0896-6273(03)00154-5

20 O’Doherty, J. P., Dayan, P., Friston, K., Critchley, H. & Dolan, R. J. Temporal difference models and reward-related learning in the human brain. Neuron 38, 329–337 (2003). 10.1016/s0896-6273(03)00169-7

21 Rutledge, R. B., Dean, M., Caplin, A. & Glimcher, P. W. Testing the Reward Prediction Error Hypothesis with an Axiomatic Model. The Journal of Neuroscience 30, 13525–13536 (2010). 10.1523/jneurosci.1747-10.2010

22 Seymour, B., Daw, N., Dayan, P., Singer, T. & Dolan, R. Differential Encoding of Losses and Gains in the Human Striatum. The Journal of Neuroscience 27, 4826–4831 (2007). 10.1523/jneurosci.0400-07.2007

23 Tobler, P. N., O’Doherty, J. P., Dolan, R. J. & Schultz, W. Human Neural Learning Depends on Reward Prediction Errors in the Blocking Paradigm. Journal of Neurophysiology 95, 301–310 (2006). 10.1152/jn.00762.2005

24 Viviani, R. et al. Effects of genetic variability of CYP2D6 on neural substrates of sustained attention during on-task activity. Translational Psychiatry 10 (2020). 10.1038/s41398-020-01020-z

25 Sarter, M., Gehring, W. J. & Kozak, R. More attention must be paid: The neurobiology of attentional effort. Brain Research Reviews 51, 145–160 (2006). 10.1016/j.brainresrev.2005.11.002

26 Shenhav, A. et al. Toward a Rational and Mechanistic Account of Mental Effort. Annual Review of Neuroscience 40, 99–124 (2017). 10.1146/annurev-neuro-072116-031526

27 Kool, W., McGuire, J. T., Rosen, Z. B. & Botvinick, M. M. Decision making and the avoidance of cognitive demand. J Exp Psychol Gen 139, 665–682 (2010). 10.1037/a0020198

28 Mcguire, J. T. & Botvinick, M. M. Prefrontal cortex, cognitive control, and the registration of decision costs. Proceedings of the National Academy of Sciences 107, 7922–7926 (2010). 10.1073/pnas.0910662107

29 Botvinick, M. & Braver, T. Motivation and Cognitive Control: From Behavior to Neural Mechanism. Annual Review of Psychology 66, 83–113 (2015). 10.1146/annurev-psych-010814-015044

30 Boksem, M. A. S. & Tops, M. Mental fatigue: Costs and benefits. Brain Research Reviews 59, 125–139 (2008). 10.1016/j.brainresrev.2008.07.001

31 Walton, M. E., Kennerley, S. W., Bannerman, D. M., Phillips, P. E. M. & Rushworth, M. F. S. Weighing up the benefits of work: Behavioral and neural analyses of effort-related decision making. Neural Networks 19, 1302–1314 (2006). 10.1016/j.neunet.2006.03.005

32 Shenhav, A., Cohen, J. D. & Botvinick, M. M. Dorsal anterior cingulate cortex and the value of control. Nature Neuroscience 19, 1286–1291 (2016). 10.1038/nn.4384

33 Salamone, J. D., Correa, M., Farrar, A. & Mingote, S. M. Effort-related functions of nucleus accumbens dopamine and associated forebrain circuits. Psychopharmacology 191, 461–482 (2007). 10.1007/s00213-006-0668-9

34 Kurzban, R., Duckworth, A., Kable, J. W. & Myers, J. An opportunity cost model of subjective effort and task performance. Behavioral and Brain Sciences 36, 661–679 (2013). 10.1017/s0140525x12003196

35 Kurzban, R. The sense of effort. Current Opinion in Psychology 7, 67–70 (2016). 10.1016/j.copsyc.2015.08.003

36 Botvinick, M. M., Huffstetler, S. & McGuire, J. T. Effort discounting in human nucleus accumbens. *Cognitive, Affective*, & Behavioral Neuroscience 9, 16–27 (2009). 10.3758/CABN.9.1.16

37 Kurniawan, I. T., Guitart-Masip, M., Dayan, P. & Dolan, R. J. Effort and Valuation in the Brain: The Effects of Anticipation and Execution. The Journal of Neuroscience 33, 6160–6169 (2013). 10.1523/jneurosci.4777-12.2013

38 Walton, M. E. & Bouret, S. What Is the Relationship between Dopamine and Effort? Trends in Neurosciences 42, 79–91 (2019). 10.1016/j.tins.2018.10.001

39 Aston-Jones, G., Rajkowski, J., Kubiak, P. & Alexinsky, T. Locus coeruleus neurons in monkey are selectively activated by attended cues in a vigilance task. The Journal of Neuroscience 14, 4467–4480 (1994).

40 Usher, M., Cohen, J. D., Servan-Schreiber, D., Rajkowski, J. & Aston-Jones, G. The Role of Locus Coeruleus in the Regulation of Cognitive Performance. Science 283, 549–554 (1999). 10.1126/science.283.5401.549

41 Mesulam, M. M. Cholinergic circuitry of the human nucleus basalis and its fate in Alzheimer’s disease. Journal of Comparative Neurology 521, 4124–4144 (2013). 10.1002/cne.23415

42 Mesulam, M. M., Mufson, E. J., Levey, A. I. & Wainer, B. H. Cholinergic innervation of cortex by the basal forebrain: cytochemistry and cortical connections of the septal area, diagonal band nuclei, nucleus basalis (substantia innominata), and hypothalamus in the rhesus monkey. J Comp Neurol 214, 170–197 (1983). 10.1002/cne.902140206

43. Mesulam, M. M. The cholinergic innervation of the human cerebral cortex in Progress in Brain Research Vol. 145 67–78 (Elsevier, 2004).

44 Broussard, J. I. Posterior parietal cortex dynamically ranks topographic signals via cholinergic influence. Front Integr Neurosci 6, 32 (2012). 10.3389/fnint.2012.00032

45. Sarter, M. & Bruno, J. P.Vigilance in Encyclopedia of the Human Brain (ed V. S. Ramachandran) 687–699 (Academic Press, 2002).

46 Sarter, M., Givens, B. & Bruno, J. P. The cognitive neuroscience of sustained attention: where top-down meets bottom-up. Brain Research Reviews 35, 146–160 (2001). 10.1016/S0165-0173(01)00044-3

47 Hangya, B., Ranade, P., Sachin Lorenc, M. & Kepecs, A. Central Cholinergic Neurons Are Rapidly Recruited by Reinforcement Feedback. Cell 162, 1155–1168 (2015). 10.1016/j.cell.2015.07.057

48 Richardson, R. T. & DeLong, M. R. Nucleus basalis of Meynert neuronal activity during a delayed response task in monkey. Brain Research 399, 364–368 (1986). 10.1016/0006-8993(86)91529-5

49 Paus, T. et al. Time-related changes in neural systems underlying attention and arousal during the performance of an auditory vigilance task. J Cogn Neurosci 9, 392–408 (1997). 10.1162/jocn.1997.9.3.392

50 Coull, J. T., Frackowiak, R. S. J. & Frith, C. D. Monitoring for target objects: activation of right frontal and parietal cortices with increasing time on task. Neuropsychologia 36, 1325–1334 (1998). 10.1016/S0028-3932(98)00035-9

51 Massar, S. A., Lim, J., Sasmita, K. & Chee, M. W. Rewards boost sustained attention through higher effort: A value-based decision making approach. Biol Psychol 120, 21–27 (2016). 10.1016/j.biopsycho.2016.07.019

52 Esterman, M., Reagan, A., Liu, G., Turner, C. & DeGutis, J. Reward reveals dissociable aspects of sustained attention. J Exp Psychol Gen 143, 2287–2295 (2014). 10.1037/xge0000019

53. Norman, D. A. & Shallice, T. Attention to Action: Willed and automatic control of behavior in Consciousness and Self-Regulation: Advances in Research and Theory Volume 4 (eds Richard J. Davidson, Gary E. Schwartz, & David Shapiro) 1–18 (Springer US, 1986).

54 Shallice, T. & Burgess, P. The domain of supervisory processes and temporal organization of behaviour. Philos Trans R Soc Lond B Biol Sci 351, 1405–1411; discussion 1411-1402 (1996). 10.1098/rstb.1996.0124

55 Riccio, C. A., Reynolds, C. R., Lowe, P. & Moore, J. J. The continuous performance test: a window on the neural substrates for attention? Archives of Clinical Neuropsychology 17, 235–272 (2002). 10.1016/S0887-6177(01)00111-1

56 Dutra, S. J., Marx, B. P., Mcglinchey, R., Degutis, J. & Esterman, M. Reward Ameliorates Posttraumatic Stress Disorder-Related Impairment in Sustained Attention. Chronic Stress 2, 247054701881240 (2018). 10.1177/2470547018812400

57 Esterman, M., Poole, V., Liu, G. & Degutis, J. Modulating Reward Induces Differential Neurocognitive Approaches to Sustained Attention. Cerebral Cortex 27, 4022–4032 (2017). 10.1093/cercor/bhw214

58 Tomporowski, P. D. & Tinsley, V. F. Effects of memory demand and motivation on sustained attention in young and older adults. Am J Psychol 109, 187–204 (1996).

59 Shenhav, A., Botvinick, M., Matthew & Cohen, D., Jonathan. The Expected Value of Control: An Integrative Theory of Anterior Cingulate Cortex Function. Neuron 79, 217–240 (2013). 10.1016/j.neuron.2013.07.007

60 Barron, H. C., Garvert, M. M. & Behrens, T. E. J. Repetition suppression: a means to index neural representations using BOLD? Philosophical Transactions of the Royal Society B: Biological Sciences 371, 20150355 (2016). 10.1098/rstb.2015.0355

61 Henson, R. N. A. Neuroimaging studies of priming. Progress in Neurobiology 70, 53–81 (2003). 10.1016/S0301-0082(03)00086-8

62 Schacter, D. L. & Buckner, R. L. Priming and the Brain. Neuron 20, 185–195 (1998). 10.1016/s0896-6273(00)80448-1

63 Grill-Spector, K., Henson, R. & Martin, A. Repetition and the brain: neural models of stimulus-specific effects. Trends in Cognitive Sciences 10, 14–23 (2006). 10.1016/j.tics.2005.11.006

64 Wiggs, C. L. & Martin, A. Properties and mechanisms of perceptual priming. Current Opinion in Neurobiology 8, 227–233 (1998). 10.1016/s0959-4388(98)80144-x

65 Race, E. A., Shanker, S. & Wagner, A. D. Neural Priming in Human Frontal Cortex: Multiple Forms of Learning Reduce Demands on the Prefrontal Executive System. Journal of Cognitive Neuroscience 21, 1766–1781 (2009). 10.1162/jocn.2009.21132

66 Schacter, D. L., Dobbins, I. G. & Schnyer, D. M. Specificity of priming: a cognitive neuroscience perspective. Nature Reviews Neuroscience 5, 853–862 (2004). 10.1038/nrn1534

67 Viviani, R., Dommes, L., Bosch, J. E. & Labek, K. Segregation, connectivity, and gradients of deactivation in neural correlates of evidence in social decision making. NeuroImage 223, 117339 (2020). 10.1016/j.neuroimage.2020.117339

68 Labek, K. & Viviani, R. Functional imaging of time on task and habituation in passive exposure to faces with emotional expression. NeuroReport 36, 135–139 (2025). 10.1097/wnr.0000000000002130

69 Salamone, D., John & Correa, M. The Mysterious Motivational Functions of Mesolimbic Dopamine. Neuron 76, 470–485 (2012). 10.1016/j.neuron.2012.10.021

70 Morales, M. & Margolis, E. B. Ventral tegmental area: cellular heterogeneity, connectivity and behaviour. Nature Reviews Neuroscience 18, 73–85 (2017). 10.1038/nrn.2016.165

71 Bouarab, C., Thompson, B. & Polter, A. M. VTA GABA Neurons at the Interface of Stress and Reward. Frontiers in Neural Circuits 13 (2019). 10.3389/fncir.2019.00078

72 Creed, M. C., Ntamati, N. R. & Tan, K. R. VTA GABA neurons modulate specific learning behaviors through the control of dopamine and cholinergic systems. Frontiers in Behavioral Neuroscience 8 (2014). 10.3389/fnbeh.2014.00008

73 Eshel, N. et al. Arithmetic and local circuitry underlying dopamine prediction errors. Nature 525, 243–246 (2015). 10.1038/nature14855

74 Zhou, W.-L. et al. Activity of a direct VTA to ventral pallidum GABA pathway encodes unconditioned reward value and sustains motivation for reward. Science Advances 8, eabm5217 (2022). 10.1126/sciadv.abm5217

75 Brown, M. T. C. et al. Ventral tegmental area GABA projections pause accumbal cholinergic interneurons to enhance associative learning. Nature 492, 452–456 (2012). 10.1038/nature11657

76 Al-Hasani, R. et al. Ventral tegmental area GABAergic inhibition of cholinergic interneurons in the ventral nucleus accumbens shell promotes reward reinforcement. Nature Neuroscience 24, 1414–1428 (2021). 10.1038/s41593-021-00898-2

77 Bakhurin, K. et al. Dopamine dynamics during stimulus-reward learning in mice can be explained by performance rather than learning. Nat Commun 16, 9081 (2025). 10.1038/s41467-025-64132-4

78 Sara, Susan J. & Bouret, S. Orienting and Reorienting: The Locus Coeruleus Mediates Cognition through Arousal. Neuron 76, 130–141 (2012). 10.1016/j.neuron.2012.09.011

79 Gritton, H. J. et al. Cortical cholinergic signaling controls the detection of cues. Proceedings of the National Academy of Sciences 113, E1089–E1097 (2016). 10.1073/pnas.1516134113

80 Harati, H., Barbelivien, A., Cosquer, B., Majchrzak, M. & Cassel, J. C. Selective cholinergic lesions in the rat nucleus basalis magnocellularis with limited damage in the medial septum specifically alter attention performance in the five-choice serial reaction time task. Neuroscience 153, 72–83 (2008). 10.1016/j.neuroscience.2008.01.031

81 Liu, R. et al. Intermittent stimulation in the nucleus basalis of meynert improves sustained attention in rhesus monkeys. Neuropharmacology 137, 202–210 (2018). 10.1016/j.neuropharm.2018.04.026

82 Martinez, V. & Sarter, M. Lateralized attentional functions of cortical cholinergic inputs. Behav Neurosci 118, 984–991 (2004). 10.1037/0735-7044.118.5.984

83 McGaughy, J., Dalley, J. W., Morrison, C. H., Everitt, B. J. & Robbins, T. W. Selective Behavioral and Neurochemical Effects of Cholinergic Lesions Produced by Intrabasalis Infusions of 192 IgG-Saporin on Attentional Performance in a Five-Choice Serial Reaction Time Task. The Journal of Neuroscience 22, 1905–1913 (2002). 10.1523/jneurosci.22-05-01905.2002

84 McGaughy, J., Decker, M. W. & Sarter, M. Enhancement of sustained attention performance by the nicotinic acetylcholine receptor agonist ABT-418 in intact but not basal forebrain-lesioned rats. Psychopharmacology 144, 175–182 (1999). 10.1007/s002130050991

85 Koulousakis, P., Andrade, P., Visser-Vandewalle, V. & Sesia, T. The Nucleus Basalis of Meynert and Its Role in Deep Brain Stimulation for Cognitive Disorders: A Historical Perspective. Journal of Alzheimer’s Disease 69, 905–919 (2019). 10.3233/JAD-180133

86 Howe, W. M. et al. Prefrontal Cholinergic Mechanisms Instigating Shifts from Monitoring for Cues to Cue-Guided Performance: Converging Electrochemical and fMRI Evidence from Rats and Humans. The Journal of Neuroscience 33, 8742–8752 (2013). 10.1523/jneurosci.5809-12.2013

87 Martinez-Rubio, C., Paulk, A. C., Mcdonald, E. J., Widge, A. S. & Eskandar, E. N. Multimodal Encoding of Novelty, Reward, and Learning in the Primate Nucleus Basalis of Meynert. The Journal of Neuroscience 38, 1942–1958 (2018). 10.1523/jneurosci.2021-17.2017

88 Crouse, R. B. et al. Acetylcholine is released in the basolateral amygdala in response to predictors of reward and enhances the learning of cue-reward contingency. eLife 9, e57335 (2020). 10.7554/eLife.57335

89 Tashakori-Sabzevar, F. & Ward, R. D. Basal Forebrain Mediates Motivational Recruitment of Attention by Reward-Associated Cues. Frontiers in Neuroscience 12 (2018). 10.3389/fnins.2018.00786

90 Wolpe, N., Holton, R. & Fletcher, P. C. What Is Mental Effort: A Clinical Perspective. Biological Psychiatry 95, 1030–1037 (2024). 10.1016/j.biopsych.2024.01.022

91 Orsini, C., Huber, D. A., Bosch, J. E. & Viviani, R. Basal forebrain and neural correlates of self-regulation traits in sustained attention. bioRxiv, 2025.2008.2005.668456 (2025). 10.1101/2025.08.05.668456

92 Stöcker, T. et al. Dependence of amygdala activation on echo time: Results from olfactory fMRI experiments. NeuroImage 30, 151–159 (2006). 10.1016/j.neuroimage.2005.09.050

93 Eickhoff, S. B., Heim, S., Zilles, K. & Amunts, K. Testing anatomically specified hypotheses in functional imaging using cytoarchitectonic maps. NeuroImage 32, 570–582 (2006). 10.1016/j.neuroimage.2006.04.204

94 Eickhoff, S. B. et al. Assignment of functional activations to probabilistic cytoarchitectonic areas revisited. NeuroImage 36, 511–521 (2007). 10.1016/j.neuroimage.2007.03.060

95 Eickhoff, S. B. et al. A new SPM toolbox for combining probabilistic cytoarchitectonic maps and functional imaging data. NeuroImage 25, 1325–1335 (2005). 10.1016/j.neuroimage.2004.12.034

96 Zaborszky, L. et al. Stereotaxic probabilistic maps of the magnocellular cell groups in human basal forebrain. NeuroImage 42, 1127–1141 (2008). 10.1016/j.neuroimage.2008.05.055

97 Amunts, K. et al. Cytoarchitectonic mapping of the human amygdala, hippocampal region and entorhinal cortex: intersubject variability and probability maps. Anatomy and Embryology 210, 343–352 (2005). 10.1007/s00429-005-0025-5

98 Kedo, O. et al. Receptor-driven, multimodal mapping of the human amygdala. Brain Structure and Function 223, 1637–1666 (2018). 10.1007/s00429-017-1577-x

99 Ballard, I. C. et al. Dorsolateral Prefrontal Cortex Drives Mesolimbic Dopaminergic Regions to Initiate Motivated Behavior. Journal of Neuroscience 31, 10340–10346 (2011). 10.1523/jneurosci.0895-11.2011

100 Murty, V. P. et al. Resting state networks distinguish human ventral tegmental area from substantia nigra. NeuroImage 100, 580–589 (2014). 10.1016/j.neuroimage.2014.06.047

101 Desikan, R. S. et al. An automated labeling system for subdividing the human cerebral cortex on MRI scans into gyral based regions of interest. NeuroImage 31, 968–980 (2006). 10.1016/j.neuroimage.2006.01.021

102 Frazier, J. A. et al. Structural brain magnetic resonance imaging of limbic and thalamic volumes in pediatric bipolar disorder. Am J Psychiatry 162, 1256–1265 (2005). 10.1176/appi.ajp.162.7.1256

103 Goldstein, J. M. et al. Hypothalamic Abnormalities in Schizophrenia: Sex Effects and Genetic Vulnerability. Biological Psychiatry 61, 935–945 (2007). 10.1016/j.biopsych.2006.06.027

104 Makris, N. et al. Decreased volume of left and total anterior insular lobule in schizophrenia. Schizophrenia Research 83, 155–171 (2006). 10.1016/j.schres.2005.11.020

105 Keren, N. I., Lozar, C. T., Harris, K. C., Morgan, P. S. & Eckert, M. A. In vivo mapping of the human locus coeruleus. NeuroImage 47, 1261–1267 (2009). 10.1016/j.neuroimage.2009.06.012

106 Huber, D., Rabl, L., Orsini, C., Labek, K. & Viviani, R. The fMRI global signal and its association with the signal from cranial bone. NeuroImage 297, 120754 (2024). 10.1016/j.neuroimage.2024.120754

107 Ramsay, J. & Silverman, B. Functional Data Analysis. (Springer, 1997).

108. dplyr: A Grammar of Data Manipulation v. 1.1.4 (2023).

109. tidyr: Tidy Messy Data v. 1.3.0 (2023).

110 Bates, D., Mächler, M., Bolker, B. & Walker, S. Fitting Linear Mixed-Effects Models Using lme4. Journal of Statistical Software 67, 1–48 (2015). 10.18637/jss.v067.i01

111 Wickham, H. ggplot2: Elegant Graphics for Data Analysis. (Springer-Verlag New York, 2016).

