## Supplementary Materials for "Functional imaging of time on task and the involvement of dopaminergic and cholinergic substrates in cognitive effort and reward"

#### Supplementary R Markdown

##### Create a general dataframe

The dataframes contain information for all of the 415 participants. Data is the main dataframe.

```
RTData <- read.table('RTData.txt', header = T,  sep='\t')
data <- rename(RTData, 'switch' = 'PI')
names(data)
```

```
##  [1] "SubjectID"    "code"         "response"     "capital"      "ISI"
##  [6] "switch"       "time"         "RT"           "block"        "trialInBlock"
## [11] "trial"
```

We rescale some variables to aid convergence of lmer. We also center the interstimulus interval (ISI) times and the switch, so that zero ISI and zero switch in the first trial are neutral (there are no switch and ISI for this trial).

```
data$block <- data$block / 10
data$firstTrialInBlock <- as.numeric(data$trialInBlock == 1)
data$trialInBlock <- data$trialInBlock / 10

idx <- data$ISI > 0
meanISI <- mean(data$ISI[idx])
summary(data$ISI[idx])
```

```
##    Min. 1st Qu.  Median    Mean 3rd Qu.    Max.
##  0.7994  1.0133  1.2133  1.2283  1.4134  1.8181
```

```
data$ISI[idx] <- data$ISI[idx] - meanISI
meanSwitch <- mean(data$switch[idx])
data$switch[idx] <- data$switch[idx] - meanSwitch
rm(idx)
sum(is.na(data)) #the 207 na are in correspondence of miss/other trials (na in RT)
```

```
## [1] 207
```

### Descriptive analysis

#### Plot RT

The plot shows that 0.25s may be used as threshold for too rapid answers. Also, there are RTs at the top that are due to response times slightly longer than 0.8s.

```
ggplot(data, aes(x= trial, y=RT)) + geom_point(alpha = 0.1) + labs(title = "RT",
    x = "Trials", y = "RT (sec)") + geom_hline(yintercept = 0.25, color = "red") +
    geom_hline(yintercept = 0.8, color = "blue") + theme_minimal()
```

```
## Warning: Removed 207 rows containing missing values or values outside the scale range
## (`geom_point()`).
```

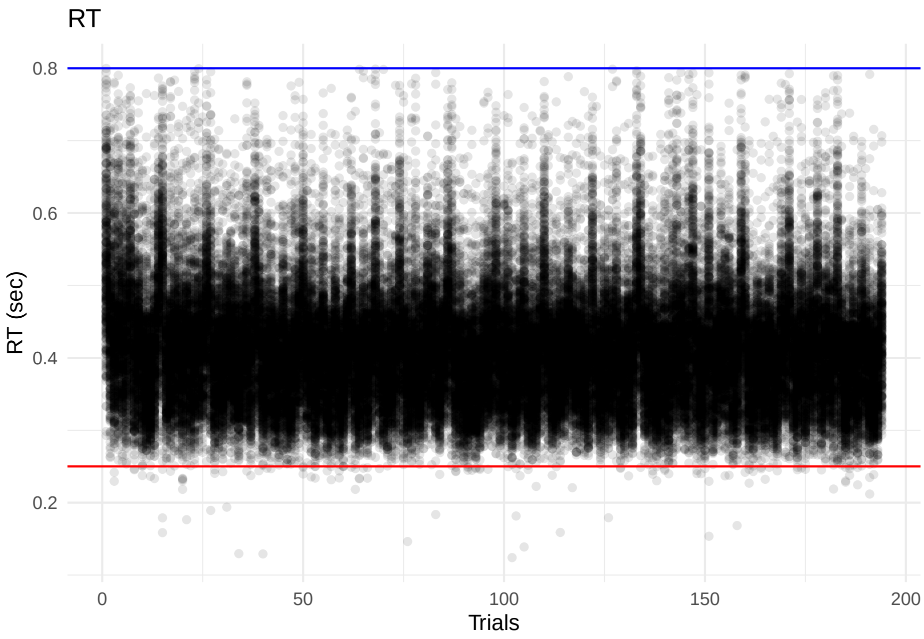


Hence, we clean up data by removing these trials. We also generate a dataframe for the analysis on accuracy.

```
dataAccuracy <- filter(data, response == "hit" | response == "incorrect" |
    response == "miss")

data <- filter(data, response == "hit" | response == "incorrect", RT > 0.25,
    RT < 0.8)
```

We examine here the response times (previously censored to be larger than 0.25s and smaller than 0.8s). The histogram shows that, when modelling, it may be more appropriate to log-transform these times.

```
par(mfrow=c(1,2))
hist(data$RT, main = "Histogram of RTs", xlab = "RTs")
hist(log(data$RT), main= "Histogram of log RTs", xlab = "log(RTs)")
```

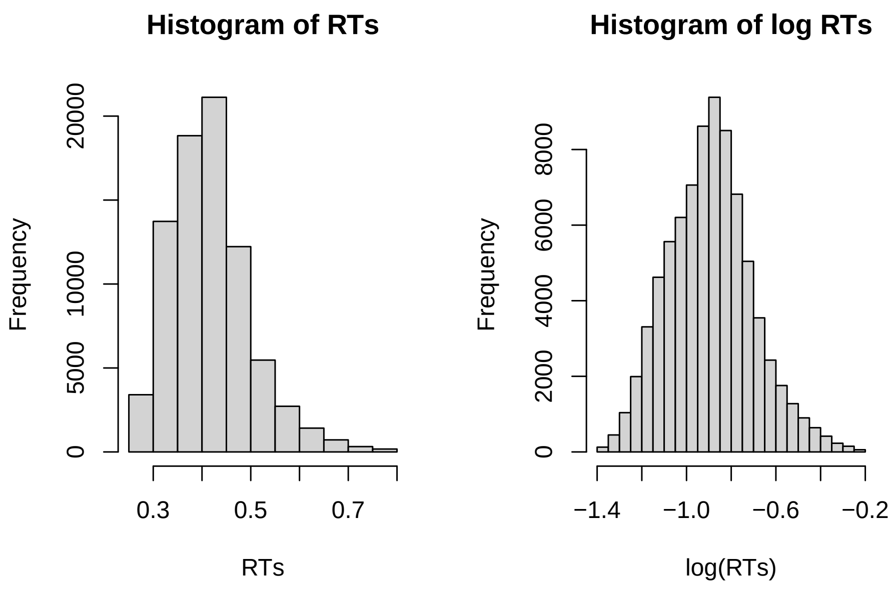


#### Dataframe for behavioral analysis

Information on sex (variable Female), Age, and acquisition site (Bonn) are added.

```
dataAgeFemale <- read.table("Query_Foraging_415_denoised_Age_Female.txt",
    sep = "\t", header = T)
dataAgeFemale$Age.z <- scale(dataAgeFemale$Age)[,1] #rescale age to aid convergence

dataBonn <- read.table("Query_Foraging_415_denoised_Bonn.txt", sep = "\t",
    header = T)

data <- left_join(data, dataBonn, by = "SubjectID") %>% left_join(dataAgeFemale,
    by = "SubjectID")

dataAccuracy <- left_join(dataAccuracy, dataBonn, by = "SubjectID") %>%
    left_join(dataAgeFemale, by = "SubjectID")
```

##### Extract demographic information

```
sum(dataAgeFemale$Female == 1)
```

```
## [1] 234
```

```
summary(dataAgeFemale$Age)
```

```
##    Min. 1st Qu.  Median    Mean 3rd Qu.    Max.
##   18.00   21.00   23.00   23.43   25.00   45.00
```

```r
sd(dataAgeFemale$Age)
```

```
## [1] 3.804921
```

```r
sum(dataBonn$Bonn == 1)
```

```
## [1] 58
```

```r
rm(dataAgeFemale)
rm(dataBonn)
```

#### RTs

##### Time on task effects (response times within the foraging block)

The following model of response times shows the first response to be much slower. The trials prosecution within block has a significant and positive effect on RTs, which means that participants get slower with increasing time on task. It turns out, however, that there are very important predictors of response time such as the ISI, and whether there was a switch in the position of the target over successive trials:

```r
fitbase <- lmer(log(RT) ~  Bonn + firstTrialInBlock + trialInBlock + switch +
    ISI + block + code + (1 | SubjectID), data = data)
summary(fitbase)
```

```
## Linear mixed model fit by REML. t-tests use Satterthwaite's method [
## lmerModLmerTest]
## Formula: log(RT) ~ Bonn + firstTrialInBlock + trialInBlock + switch +
##     ISI + block + code + (1 | SubjectID)
##    Data: data
##
## REML criterion at convergence: -125997.4
##
## Scaled residuals:
##     Min      1Q  Median      3Q     Max
## -4.5199 -0.6367 -0.0894  0.5219  6.7638
##
## Random effects:
##  Groups    Name        Variance Std.Dev.
##  SubjectID (Intercept) 0.01657  0.1287
##  Residual              0.01179  0.1086
## Number of obs: 80133, groups:  SubjectID, 415
##
## Fixed effects:
##                     Estimate Std. Error         df t value Pr(>|t|)
## (Intercept)       -8.729e-01  6.936e-03  4.403e+02 -125.84  < 2e-16 ***
## Bonn              -1.612e-01  1.826e-02  4.130e+02   -8.83  < 2e-16 ***
## firstTrialInBlock  1.883e-01  1.592e-03  7.971e+04  118.24  < 2e-16 ***
## trialInBlock       6.259e-03  1.259e-03  7.971e+04    4.97 6.69e-07 ***
## switch            -6.728e-02  8.148e-04  7.971e+04  -82.57  < 2e-16 ***
## ISI               -1.275e-01  1.458e-03  7.971e+04  -87.47  < 2e-16 ***
```

```
## block             -2.450e-02  8.339e-04  7.971e+04  -29.37  < 2e-16 ***
## codelow            7.497e-03  7.714e-04  7.971e+04    9.72  < 2e-16 ***
## ---
## Signif. codes:  0 '***' 0.001 '**' 0.01 '*' 0.05 '.' 0.1 ' ' 1
##
## Correlation of Fixed Effects:
##             (Intr) Bonn   frsTIB trlInB switch ISI    block
## Bonn        -0.368
## frstTrlInBl -0.076  0.000
## trialInBlck -0.129  0.000  0.480
## switch      -0.010  0.000  0.021  0.044
## ISI          0.013  0.000 -0.053 -0.111  0.088
## block       -0.108  0.000  0.004  0.009  0.018  0.008
## codelow     -0.065  0.000  0.002  0.003  0.046  0.007  0.093
```

We can see that responses are quicker when the ISI was longer, and when the target switched sides from one to the next trial. To visualize the effect of ISI, we plot it after residualizing for unwanted clouding variance.

```
fittmp <- lmer(log(RT) ~ Bonn + trialInBlock + switch + (1 | block) +
    (1 | SubjectID), data = data, subset = firstTrialInBlock == 0)

ggplot(data[data$firstTrialInBlock == 0,], aes(x = ISI, y = resid(fittmp))) +
    geom_smooth(se = FALSE) + geom_point(alpha = 0.01) + theme_minimal() +
    ylab("log adjusted RT") + xlab("ISI (centered)")
```

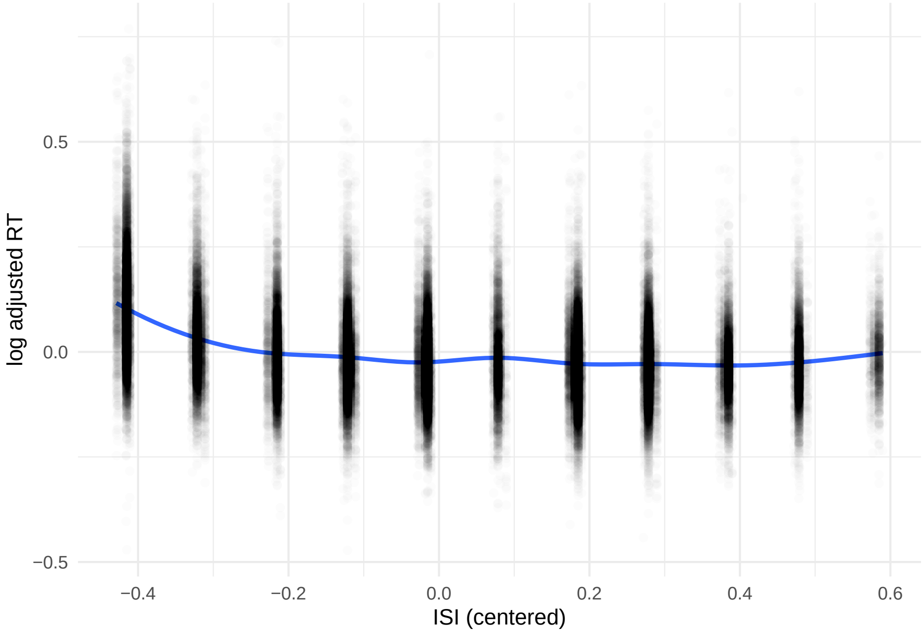


We can see that the effect of ISI plateaus rapidly at the average ISI. It is therefore appropriate to model ISI with a non-linear effect. We find that a cubic polynomial addresses this problem reasonably well.

```
ggplot(data[data$firstTrialInBlock == 0,], aes(x = ISI, y = resid(fittmp))) +
    geom_smooth(aes(color = "Loess")) + geom_smooth(aes(color = "Cubic polynomial"),
    formula = y~poly(x,3)) + scale_color_manual(name= "ISI Fit", values =
```

```r
    c("Loess" = "blue", "Cubic polynomial" = "red")) + theme_minimal() +
    ylab("log adjusted RT") + xlab("ISI (centered)")
```

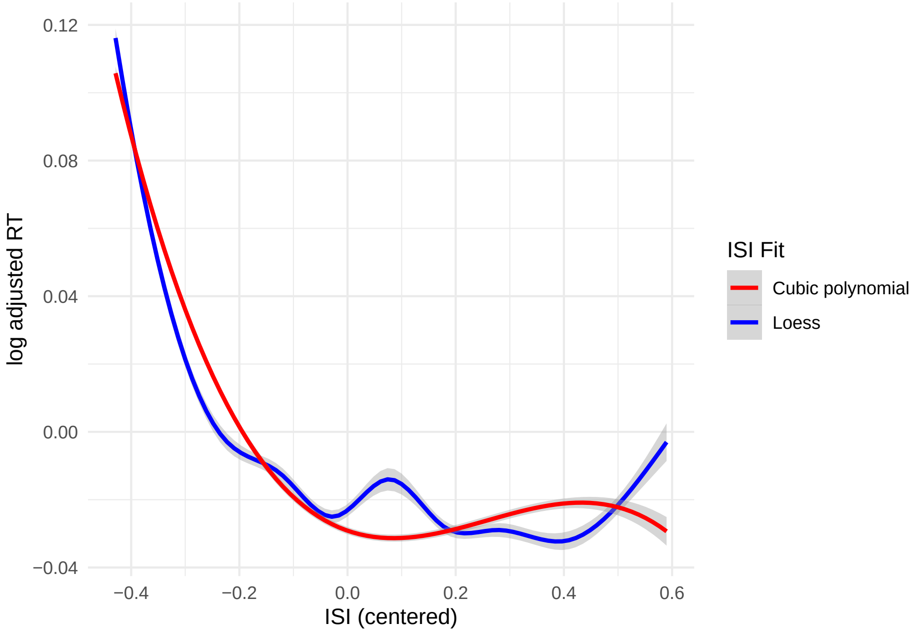


```r
rm(fittmp)
```

We now model again response times using the polynomial fit for ISI and switch as covariates.

```r
fitbase <- lmer(log(RT) ~  Bonn + firstTrialInBlock + trialInBlock + switch +
    poly(ISI,3) + block + code + (1 | SubjectID), data = data)
summary(fitbase)
```

```
## Linear mixed model fit by REML. t-tests use Satterthwaite's method [
## lmerModLmerTest]
## Formula: log(RT) ~ Bonn + firstTrialInBlock + trialInBlock + switch +
##     poly(ISI, 3) + block + code + (1 | SubjectID)
##    Data: data
##
## REML criterion at convergence: -131487.3
##
## Scaled residuals:
##     Min      1Q  Median      3Q     Max
## -5.1429 -0.6254 -0.0777  0.5178  7.2050
##
## Random effects:
##  Groups    Name        Variance Std.Dev.
##  SubjectID (Intercept) 0.01660  0.1288
##  Residual              0.01101  0.1049
## Number of obs: 80133, groups:  SubjectID, 415
##
## Fixed effects:
```

```
##                     Estimate Std. Error         df  t value Pr(>|t|)
## (Intercept)       -8.732e-01  6.934e-03  4.386e+02 -125.931  < 2e-16 ***
## Bonn              -1.614e-01  1.827e-02  4.130e+02   -8.832  < 2e-16 ***
## firstTrialInBlock  2.182e-01  1.591e-03  7.971e+04  137.174  < 2e-16 ***
## trialInBlock       4.237e-03  1.220e-03  7.971e+04    3.472 0.000516 ***
## switch            -6.926e-02  7.890e-04  7.971e+04  -87.779  < 2e-16 ***
## poly(ISI, 3)1     -9.605e+00  1.061e-01  7.971e+04  -90.563  < 2e-16 ***
## poly(ISI, 3)2      7.748e+00  1.095e-01  7.971e+04   70.776  < 2e-16 ***
## poly(ISI, 3)3     -2.917e+00  1.059e-01  7.971e+04  -27.548  < 2e-16 ***
## block             -2.465e-02  8.061e-04  7.971e+04  -30.574  < 2e-16 ***
## codelow            6.025e-03  7.455e-04  7.971e+04    8.082 6.46e-16 ***
## ---
## Signif. codes:  0 '***' 0.001 '**' 0.01 '*' 0.05 '.' 0.1 ' ' 1
##
## Correlation of Fixed Effects:
##             (Intr) Bonn   frsTIB trlInB switch p(ISI,3)1 p(ISI,3)2 p(ISI,3)3
## Bonn        -0.368
## frstTrlInBl -0.072  0.000
## trialInBlck -0.126  0.000  0.461
## switch      -0.009  0.000  0.009  0.040
## ply(ISI,3)1  0.013  0.000 -0.052 -0.112  0.088
## ply(ISI,3)2 -0.005  0.000  0.251  0.005 -0.051 -0.005
## ply(ISI,3)3 -0.012  0.000 -0.047  0.077 -0.042 -0.013    -0.023
## block       -0.105  0.000  0.004  0.012  0.016  0.007     0.009     0.030
## codelow     -0.063  0.000 -0.005  0.004  0.046  0.007    -0.022     0.016
##             block
## Bonn
## frstTrlInBl
## trialInBlck
## switch
## ply(ISI,3)1
## ply(ISI,3)2
## ply(ISI,3)3
## block
## codelow      0.093
```

We can see that response times increase with time on task, and this result does not depend on ISI modeling: also linear and quadratic effects give a significant effect of time on task.

##### Forage block effects

Forage block effects are between trials and therefore not immediately relevant for time on task effects. However, we examine them here because the decrease in response times with the progression of the blocks, when modeled linearly, suggests a different mechanism than the progression of trials within block, such as a learning effect.

We want to visualize the effect of blocks after adjusting for ISI, code etc. The effect of the trials in block is modeled non-parametrically with a random effect. The plot shows that after an initial decline, RTs oscillate at a lower level. The composition of trials is not a good explanation of these oscillations, since trials were modeled as random effects. The first trial in the block was excluded here, so it also cannot contribute to these oscillations.

```r
data_tail <- filter(data, firstTrialInBlock == 0) %>% drop_na()
fitblocks <- lmer(log(RT) ~ Bonn + switch + poly(ISI,3) + code +
    (1 | block : trialInBlock) + (1 | SubjectID), data = data_tail)
summary(fitblocks)
```

```
## Linear mixed model fit by REML. t-tests use Satterthwaite's method [
## lmerModLmerTest]
## Formula:
## log(RT) ~ Bonn + switch + poly(ISI, 3) + code + (1 | block:trialInBlock) +
##     (1 | SubjectID)
##    Data: data_tail
##
## REML criterion at convergence: -131944.9
##
## Scaled residuals:
##     Min      1Q  Median      3Q     Max
## -6.0945 -0.6160 -0.0716  0.5136  7.9824
##
## Random effects:
##  Groups             Name        Variance Std.Dev.
##  SubjectID          (Intercept) 0.016864 0.12986
##  block:trialInBlock (Intercept) 0.001122 0.03349
##  Residual                       0.009334 0.09661
## Number of obs: 73552, groups:  SubjectID, 415; block:trialInBlock, 178
##
## Fixed effects:
##                   Estimate Std. Error         df  t value Pr(>|t|)
## (Intercept)    -0.887629   0.007754 564.035043 -114.470  < 2e-16 ***
## Bonn           -0.167047   0.018414 413.001004   -9.072  < 2e-16 ***
## switch         -0.070184   0.004703 249.557693  -14.922  < 2e-16 ***
## poly(ISI, 3)1 -10.433711   0.640539 228.024969  -16.289  < 2e-16 ***
## poly(ISI, 3)2   7.045627   0.585659 325.301187   12.030  < 2e-16 ***
## poly(ISI, 3)3  -2.817909   0.616504 266.712836   -4.571 7.44e-06 ***
## codelow         0.005873   0.005077 170.381989    1.157    0.249
## ---
## Signif. codes:  0 '***' 0.001 '**' 0.01 '*' 0.05 '.' 0.1 ' ' 1
##
## Correlation of Fixed Effects:
##             (Intr) Bonn   switch p(ISI,3)1 p(ISI,3)2 p(ISI,3)3
## Bonn        -0.332
## switch      -0.014  0.000
## ply(ISI,3)1 -0.002  0.002  0.073
## ply(ISI,3)2  0.006  0.000 -0.058 -0.077
## ply(ISI,3)3 -0.004  0.001 -0.056 -0.040    -0.014
## codelow     -0.327  0.000  0.043  0.007    -0.021     0.011
```

```r
ggplot(data_tail, aes(x = block*10, y = resid(fitblocks))) +
    geom_smooth(aes(color = "Linear"), formula = y~x, se = F) +
    geom_smooth(aes(color="Cubic polynomial"), formula = y~poly(x,3), se=F) +
    geom_smooth(aes(color="Logarithm"), formula = y ~ log(x), se=F) +
    theme_minimal() + ylab("log adjusted RT") + xlab("Block") +
    scale_color_manual(name= "Blocks Fit", values= c("Linear" = "blue",
    "Cubic polynomial" = "red", "Logarithm"  = "darkgreen"))
```

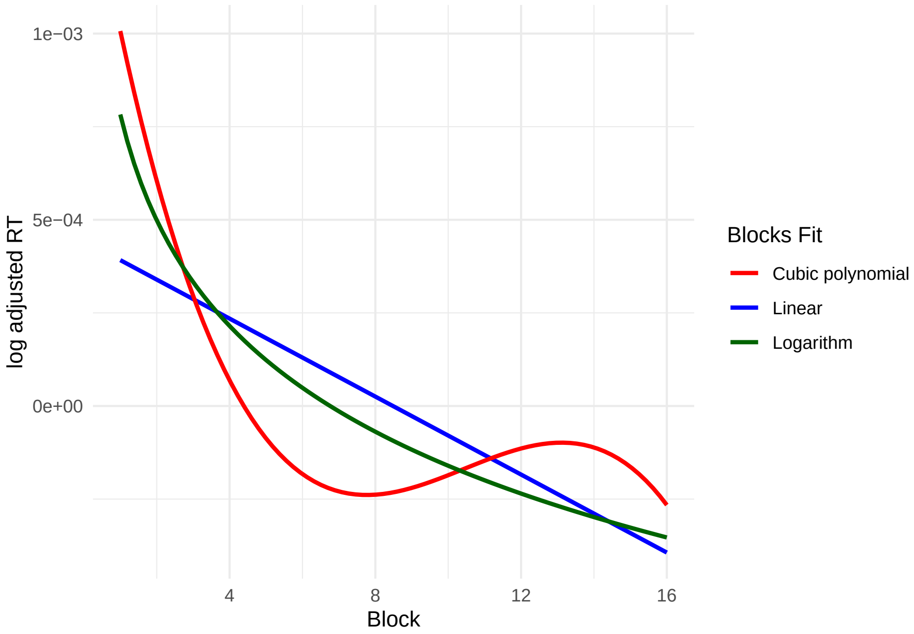


A log transform of block appears to be sufficient. Therefore, we use the log transform in the sequel.

##### Response times model

This gives the following model:

```
fitbase <- lmer(log(RT) ~ firstTrialInBlock + trialInBlock + switch + poly(ISI,3) +
    code + log(block) + Bonn + (1 | SubjectID), data = data)
summary(fitbase)
```

```
## Linear mixed model fit by REML. t-tests use Satterthwaite's method [
## lmerModLmerTest]
## Formula: log(RT) ~ firstTrialInBlock + trialInBlock + switch + poly(ISI,
##     3) + code + log(block) + Bonn + (1 | SubjectID)
##    Data: data
##
## REML criterion at convergence: -132293.3
##
## Scaled residuals:
##     Min      1Q  Median      3Q     Max
## -5.3660 -0.6243 -0.0739  0.5182  7.2535
##
## Random effects:
##  Groups    Name        Variance Std.Dev.
##  SubjectID (Intercept) 0.0166   0.1289
##  Residual              0.0109   0.1044
## Number of obs: 80133, groups:  SubjectID, 415
##
## Fixed effects:
##                       Estimate Std. Error         df  t value Pr(>|t|)
## (Intercept)        -9.002e-01  6.896e-03  4.290e+02 -130.530  < 2e-16 ***
```

```
## firstTrialInBlock  2.181e-01  1.583e-03  7.971e+04  137.806  < 2e-16 ***
## trialInBlock       3.539e-03  1.214e-03  7.971e+04    2.915  0.00356 **
## switch            -7.019e-02  7.855e-04  7.971e+04  -89.347  < 2e-16 ***
## poly(ISI, 3)1     -9.606e+00  1.055e-01  7.971e+04  -91.028  < 2e-16 ***
## poly(ISI, 3)2      7.790e+00  1.089e-01  7.971e+04   71.526  < 2e-16 ***
## poly(ISI, 3)3     -2.984e+00  1.054e-01  7.971e+04  -28.322  < 2e-16 ***
## codelow            3.282e-03  7.476e-04  7.971e+04    4.390 1.13e-05 ***
## log(block)        -2.029e-02  4.842e-04  7.971e+04  -41.897  < 2e-16 ***
## Bonn              -1.614e-01  1.827e-02  4.130e+02   -8.831  < 2e-16 ***
## ---
## Signif. codes:  0 '***' 0.001 '**' 0.01 '*' 0.05 '.' 0.1 ' ' 1
##
## Correlation of Fixed Effects:
##              (Intr) frsTIB trlInB switch p(ISI,3)1 p(ISI,3)2 p(ISI,3)3 codelw
## frstTrlInBl -0.072
## trialInBlck -0.124  0.461
## switch      -0.007  0.009  0.041
## ply(ISI,3)1  0.014 -0.052 -0.112  0.088
## ply(ISI,3)2 -0.004  0.251  0.005 -0.052 -0.005
## ply(ISI,3)3 -0.008 -0.047  0.077 -0.041 -0.013    -0.024
## codelow     -0.051 -0.004  0.006  0.051  0.007    -0.023     0.019
## log(block)   0.016  0.005  0.022  0.040  0.005    -0.003     0.038     0.155
## Bonn        -0.370  0.000  0.000  0.000  0.000     0.000     0.000     0.000
##              lg(bl)
## frstTrlInBl
## trialInBlck
## switch
## ply(ISI,3)1
## ply(ISI,3)2
## ply(ISI,3)3
## codelow
## log(block)
## Bonn         0.000
```

We look at the residuals:

```
hist(resid(fitbase), breaks = 30, main = "Histogram of residuals: adjusted log RTs",
    xlab ="Residuals (log adjusted RTS)")
```

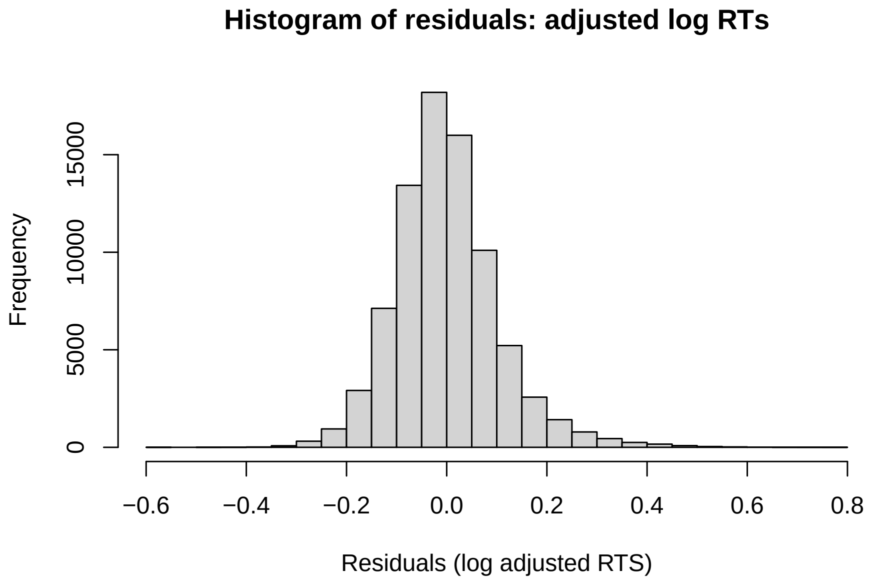


We verify that this fit models the response times by comparing the real with the fitted data:

```
data0 <- select(data, RT, trialInBlock)
data0$RTfit <- exp(fitted(fitbase) + rnorm(n=dim(data0)[1], mean=0,
    sd=sd(exp(resid(fitbase)))))
par(mfrow=c(1,2))
hist(data0$RT,  main = "Real data histogram", xlab = "RTs")
hist(data0$RTfit, main = "Fitted data histogram", xlab = "Fitted RTs")
```

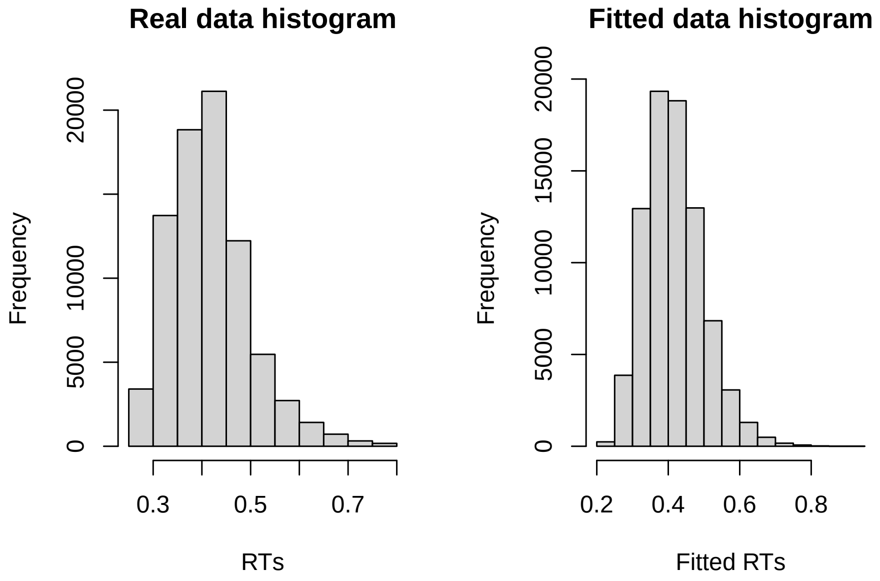

```r
rm(data0)
```

The post-fit check shows the model to assume residuals that are slightly overdispersed relative to the real data. However, the model appears to fit the data fairly well. From this model, we may say that:

- time on task is associated with a small increase in response times;
- the progression of the blocks, in contrast, with a larger decrease in response times.

We may use a simplified fit on non-logged response times to make the coefficients more easily interpretable. As one can see, from an inferential point of view, the conclusions do not differ from those of the more sophisticated model. To aid the interpretation, RTs and ISI are converted in ms. The variables trial in block and block were previously scaled by dividing them by 10 to aid convergence. Therefore, their coefficients must be divided by 10 as well to interpret their effects previous to scaling.

```r
data$ISIms <- data$ISI * 1000
summary(lmer(RT*1000 ~ firstTrialInBlock + trialInBlock + switch + ISIms + code +
    block + Bonn + (1 | SubjectID), data = data))
```

```
## Linear mixed model fit by REML. t-tests use Satterthwaite's method [
## lmerModLmerTest]
## Formula: RT * 1000 ~ firstTrialInBlock + trialInBlock + switch + ISIms +
##     code + block + Bonn + (1 | SubjectID)
##    Data: data
##
## REML criterion at convergence: 853625.5
##
## Scaled residuals:
##     Min      1Q  Median      3Q     Max
## -4.2241 -0.6095 -0.1212  0.4413  8.0248
##
## Random effects:
##  Groups    Name        Variance Std.Dev.
##  SubjectID (Intercept) 2993     54.71
##  Residual              2408     49.07
## Number of obs: 80133, groups:  SubjectID, 415
##
## Fixed effects:
##                     Estimate Std. Error         df t value Pr(>|t|)
## (Intercept)        4.250e+02  2.954e+00  4.439e+02 143.865  < 2e-16 ***
## firstTrialInBlock  8.455e+01  7.195e-01  7.971e+04 117.505  < 2e-16 ***
## trialInBlock       2.536e+00  5.690e-01  7.971e+04   4.458 8.30e-06 ***
## switch            -2.737e+01  3.681e-01  7.971e+04 -74.340  < 2e-16 ***
## ISIms             -5.655e-02  6.586e-04  7.971e+04 -85.861  < 2e-16 ***
## codelow            3.160e+00  3.485e-01  7.971e+04   9.066  < 2e-16 ***
## block             -1.101e+01  3.768e-01  7.971e+04 -29.218  < 2e-16 ***
## Bonn              -6.304e+01  7.761e+00  4.130e+02  -8.122 5.34e-15 ***
## ---
## Signif. codes:  0 '***' 0.001 '**' 0.01 '*' 0.05 '.' 0.1 ' ' 1
##
## Correlation of Fixed Effects:
##             (Intr) frsTIB trlInB switch ISIms  codelw block
## frstTrlInBl -0.081
```

```
## trialInBlck -0.137  0.480
## switch      -0.011  0.021  0.044
## ISIms        0.014 -0.053 -0.111  0.088
## codelow     -0.069  0.002  0.003  0.046  0.007
## block       -0.115  0.004  0.009  0.018  0.008  0.093
## Bonn        -0.367  0.000  0.000  0.000  0.000  0.000  0.000
```

We see here that an increase of 100 msec in the ISI leads to a reduction in reaction times of 5.7 msec, while a switch of target to the contralateral side to a reduction of 27.4 msec (both $p < 0.001$). Participants were quicker in the high reward condition of about 3.2 msec ($p < 0.001$). The progression of trials leads to an increase in reaction times of about 0.3 msec (2.5/10) per trial ($t = 4.5$, $p < 0.001$). This increase is consistent with the time on task during the forage path constituting a cognitive challenge for participants, and it is aligned with the literature on sustained attention. Note that the progression of blocks, in contrast, led to an average reduction of reaction times of about 1.1 msec (-11.01/10) per block ($p < 0.001$). In reality, as previously shown, this is best modeled by a nonlinear effect, because the first blocks are more effective in reducing reaction times than the last. Hence, it is the time on task within forage block that represents the cognitive challenge, not the block prosecution.

#### Accuracy

When excluding the response category "other" (therefore including misses and incorrect responses), global accuracy is over the 99%.

```
summary(dataAccuracy$response) #80445 total hit/miss/incorrect responses
```

```
##    Length     Class      Mode
##     80445 character character
```

```
summary(dataAccuracy$response == "hit")
```

```
##    Mode   FALSE    TRUE
## logical     402   80043
```

```
accuracy_percent <- (80043/80445)*100
accuracy_percent
```

```
## [1] 99.50028
```

To avoid convergence issues in the accuracy models, we center all the predictors excluded code. ISI and switch have been previously centered.

```
dataAccuracy$firstTrialInBlock_centered <- scale(dataAccuracy$firstTrialInBlock,
    center = T, scale = F)
dataAccuracy$Bonn_centered <- scale(dataAccuracy$Bonn, center = T, scale = F)
dataAccuracy$trialInBlock_centered <- scale(dataAccuracy$trialInBlock, center = T,
    scale = F)
dataAccuracy$block_centered <- scale(dataAccuracy$block, center = T, scale = F)

summary((glmer(response == "hit" ~ firstTrialInBlock_centered +
    trialInBlock_centered + switch + ISI + code + block_centered + Bonn_centered +
    (1 | SubjectID), data = dataAccuracy, family = binomial)))
```

```
## Generalized linear mixed model fit by maximum likelihood (Laplace
##   Approximation) [glmerMod]
##  Family: binomial  ( logit )
## Formula:
## response == "hit" ~ firstTrialInBlock_centered + trialInBlock_centered +
##     switch + ISI + code + block_centered + Bonn_centered + (1 |      SubjectID)
##    Data: dataAccuracy
##
##      AIC      BIC   logLik deviance df.resid
##   4746.4   4830.1  -2364.2   4728.4    80436
##
## Scaled residuals:
##     Min      1Q  Median      3Q     Max
## -32.080   0.035   0.052   0.072   0.384
##
## Random effects:
##  Groups    Name        Variance Std.Dev.
##  SubjectID (Intercept) 0.8671   0.9312
## Number of obs: 80445, groups:  SubjectID, 415
##
## Fixed effects:
##                            Estimate Std. Error z value Pr(>|z|)
## (Intercept)                  6.1890     0.1160  53.340  < 2e-16 ***
## firstTrialInBlock_centered  -1.4635     0.1852  -7.902 2.75e-15 ***
## trialInBlock_centered       -0.7349     0.1749  -4.201 2.66e-05 ***
## switch                       1.3071     0.1241  10.530  < 2e-16 ***
## ISI                          0.7038     0.2038   3.454 0.000553 ***
## codelow                     -0.3661     0.1027  -3.567 0.000362 ***
## block_centered               0.1388     0.1090   1.273 0.202997
## Bonn_centered                0.2795     0.2222   1.258 0.208551
## ---
## Signif. codes:  0 '***' 0.001 '**' 0.01 '*' 0.05 '.' 0.1 ' ' 1
##
## Correlation of Fixed Effects:
##             (Intr) frTIB_ trlIB_ switch ISI    codelw blck_c
## frstTrlInB_ -0.205
## trlInBlck_c -0.111  0.641
## switch       0.288 -0.181  0.035
## ISI          0.084 -0.112 -0.099  0.093
## codelow     -0.527  0.012  0.017  0.026  0.004
## block_cntrd  0.035 -0.007 -0.003  0.056  0.063  0.010
## Bonn_centrd  0.028  0.000  0.000  0.001  0.003  0.000  0.000
```

The main positive predictor of accuracy is the switch, as the participants were more precise when the target switched its position in consecutive trials ($z = 10.5$, $p < 0.001$). The stronger negative predictor is the first trial in block, where participants are less accurate ($z = -7.9$, $p < 0.001$). Moreover, they got less accurate with the trials in block prosecution ($z = -4.2$, $p < 0.001$) and in the low reward condition ($z = -3.6$, $p < 0.001$). On the opposite, participants were more accurate with a longer ISI ($z = 3.5$, $p < 0.001$).

#### Supplementary Results Tables

**Table S1. Foraging trend. Effect of time on task, positive direction.**

| Clu # | Brain area | Coord. (mm.) | *t* | *p* (uncorr.) | *p* (corr.) | *k* | *p* (cl.) |
|---|---|---|---|---|---|---|---|
| 1 | R Posterior Cingulate Gyrus (BA 23) | 9 -40 41 | 22.046 | < 0.0001 | <0.001 | 291531 | <0.001 |
| | R Superior Frontal Gyrus (BA 9) | 25 35 38 | 20.598 | < 0.0001 | <0.001 | | |
| | L Anterior Cingulate (BA 11) | -7 33 0 | 20.238 | < 0.0001 | <0.001 | | |
| | R Inferior Frontal Gyrus (BA 47) | 28 23 -7 | 20.143 | < 0.0001 | <0.001 | | |
| | R Superior Temporal Gyrus (BA 41) | 54 -20 5 | 19.707 | < 0.0001 | <0.001 | | |
| | R Superior Frontal Gyrus (BA 8) | 27 23 54 | 19.699 | < 0.0001 | <0.001 | | |
| | R Lentiform Nucleus (Putamen) | 31 -19 -1 | 19.387 | < 0.0001 | <0.001 | | |
| | R Precuneus (BA 31) | 9 -64 23 | 19.366 | < 0.0001 | <0.001 | | |
| | R Inferior Frontal Gyrus (BA 47) | 30 29 -16 | 19.350 | < 0.0001 | <0.001 | | |
| | R Anterior Cingulate (BA 32) | 9 33 21 | 19.335 | < 0.0001 | <0.001 | | |
| | L Insula (BA 13) | -28 18 -7 | 19.318 | < 0.0001 | <0.001 | | |
| | R Anterior Cingulate (BA 32) | 4 36 27 | 19.103 | < 0.0001 | <0.001 | | |
| | R Medial Frontal Gyrus (BA 8) | 4 26 44 | 19.102 | < 0.0001 | <0.001 | | |
| | R Lingual Gyrus (BA 18) | 22 -88 -5 | 18.690 | < 0.0001 | <0.001 | | |
| | R Anterior Cingulate (BA 11) | 9 38 5 | 18.630 | < 0.0001 | <0.001 | | |
| | R Precentral Gyrus (BA 6) | 39 -11 35 | 18.496 | < 0.0001 | <0.001 | | |
| | R Precuneus (BA 39) | 42 -70 35 | 18.297 | < 0.0001 | <0.001 | | |
| | L Caudate (Caudate Head) | -12 15 2 | 18.265 | < 0.0001 | <0.001 | | |
| | L Superior Temporal Gyrus (BA 41) | -45 -26 3 | 18.238 | < 0.0001 | <0.001 | | |
| | L Precentral Gyrus (BA 4) | -37 -14 35 | 18.096 | < 0.0001 | <0.001 | | |
| | R Inferior Parietal Lobule (BA 39) | 37 -62 42 | 17.711 | < 0.0001 | <0.001 | | |
| | R Middle Frontal Gyrus (BA 9) | 46 27 26 | 17.627 | < 0.0001 | <0.001 | | |
| | R Cuneus (BA 18) | 10 -79 23 | 17.472 | < 0.0001 | <0.001 | | |
| | L Medial Frontal Gyrus (BA 4) | -6 -26 63 | 17.419 | < 0.0001 | <0.001 | | |
| | R Inferior Frontal Gyrus, Orbital Part (BA 47) | 40 33 -17 | 17.381 | < 0.0001 | <0.001 | | |
| | L Medial Frontal Gyrus (BA 9) | -24 36 33 | 17.342 | < 0.0001 | <0.001 | | |
| | R Middle Frontal Gyrus (BA 46) | 48 32 18 | 17.231 | < 0.0001 | <0.001 | | |
| | L Precuneus (BA 23) | -7 -67 23 | 17.197 | < 0.0001 | <0.001 | | |
| | R Medial Frontal Gyrus (BA 4) | 6 -22 65 | 17.148 | < 0.0001 | <0.001 | | |
| | R Precuneus (BA 7) | 13 -77 33 | 17.077 | < 0.0001 | <0.001 | | |
| | L Cingulate Gyrus (BA 31) | -6 -38 41 | 17.057 | < 0.0001 | <0.001 | | |
| | L Inferior Frontal Gyrus (BA 47) | -28 24 -14 | 16.921 | < 0.0001 | <0.001 | | |
| | L Angular Gyrus (BA 39) | -40 -77 32 | 16.814 | < 0.0001 | <0.001 | | |
| | R Middle Temporal Gyrus (BA 21) | 60 -46 2 | 16.798 | < 0.0001 | <0.001 | | |

| | | | | |
|---|---|---|---|---|
| R Inferior Frontal Gyrus (BA 38) | 43 27 -13 | 16.781 | < 0.0001 | <0.001 |
| L Middle Frontal Gyrus (BA 8) | -24 14 51 | 16.678 | < 0.0001 | <0.001 |
| R Middle Frontal Gyrus (BA 44) | 39 12 42 | 16.512 | < 0.0001 | <0.001 |
| L Superior Frontal Gyrus (BA 8) | -25 17 57 | 16.319 | < 0.0001 | <0.001 |
| R Caudate | 16 -2 18 | 15.901 | < 0.0001 | <0.001 |
| R Lentiform Nucleus (Putamen) | 28 0 -7 | 15.843 | < 0.0001 | <0.001 |
| R Superior Temporal Gyrus (BA 22) | 52 -49 20 | 15.461 | < 0.0001 | <0.001 |
| L Precuneus (BA 19) | -36 -74 39 | 15.391 | < 0.0001 | <0.001 |
| R Superior Frontal Gyrus (BA 10) | 25 54 8 | 15.332 | < 0.0001 | <0.001 |
| R Medial Frontal Gyrus (BA 11) | 7 48 -13 | 15.210 | < 0.0001 | <0.001 |
| R Caudate (Caudate Head) | 10 14 5 | 15.159 | < 0.0001 | <0.001 |
| L Inferior Parietal Lobule (BA 7) | -31 -61 41 | 15.159 | < 0.0001 | <0.001 |
| R Medial Frontal Gyrus (BA 10) | 6 47 -10 | 15.139 | < 0.0001 | <0.001 |
| R Cuneus (BA 18) | 7 -68 3 | 15.053 | < 0.0001 | <0.001 |
| L Inferior Frontal Gyrus, Orbital Part (BA 47) | -43 32 -19 | 14.968 | < 0.0001 | <0.001 |
| L Superior Temporal Gyrus (BA 22) | -58 -29 5 | 14.829 | < 0.0001 | <0.001 |
| L Middle Frontal Gyrus (BA 46) | -48 20 24 | 14.761 | < 0.0001 | <0.001 |
| L Superior Frontal Gyrus (BA 10) | -22 53 11 | 14.538 | < 0.0001 | <0.001 |
| L Middle Temporal Gyrus (BA 21) | -63 -49 5 | 14.500 | < 0.0001 | <0.001 |
| R Middle Temporal Gyrus (BA 21) | 48 -28 -7 | 14.400 | < 0.0001 | <0.001 |
| R Anterior Prefrontal Cortex (BA 46) | 25 45 17 | 14.391 | < 0.0001 | <0.001 |
| R Superior Frontal Gyrus, Orbital Part (BA 11) | 19 54 -7 | 14.163 | < 0.0001 | <0.001 |
| L Precuneus (BA 7) | -6 -65 45 | 14.132 | < 0.0001 | <0.001 |
| L Cuneus (BA 18) | -6 -80 23 | 14.005 | < 0.0001 | <0.001 |
| R Middle Temporal Gyrus (BA 20) | 49 -19 -13 | 13.992 | < 0.0001 | <0.001 |
| R Parahippocampal Gyrus (BA 28) | 21 -13 -22 | 13.989 | < 0.0001 | <0.001 |
| L Middle Frontal Gyrus (BA 44) | -43 14 32 | 13.954 | < 0.0001 | <0.001 |
| R Middle Temporal Gyrus (BA 21) | 58 -19 -13 | 13.930 | < 0.0001 | <0.001 |
| R Superior Frontal Gyrus (BA 11) | 22 51 -1 | 13.661 | < 0.0001 | <0.001 |
| R Lingual Gyrus (BA 18) | 16 -70 -4 | 13.657 | < 0.0001 | <0.001 |
| R Inferior Parietal Lobule (BA 40) | 48 -43 50 | 13.502 | < 0.0001 | <0.001 |
| L Supramarginal Gyrus (BA 22) | -60 -52 21 | 13.066 | < 0.0001 | <0.001 |
| L Parahippocampal Gyrus (BA 28) | -21 -14 -22 | 12.959 | < 0.0001 | <0.001 |
| R Parahippocampal Gyrus (BA 19) | 19 -46 -7 | 12.887 | < 0.0001 | <0.001 |
| L Thalamus | -6 -7 -1 | 12.658 | < 0.0001 | <0.001 |
| L Cingulate Gyrus | -7 -20 29 | 12.569 | < 0.0001 | <0.001 |
| R Postcentral Gyrus (BA 2) | 61 -23 36 | 12.221 | < 0.0001 | <0.001 |
| R Thalamus | 6 -8 2 | 12.046 | < 0.0001 | <0.001 |

| | | | | | | | |
|---|---|---|---|---|---|---|---|
| | R Middle Frontal Gyrus (BA 10) | 43 54 -13 | 11.924 | < 0.0001 | <0.001 | | |
| | L Middle Frontal Gyrus (BA 47) | -34 36 -1 | 11.918 | < 0.0001 | <0.001 | | |
| | L Middle Temporal Gyrus (BA 20) | -51 -14 -14 | 11.885 | < 0.0001 | <0.001 | | |
| | R Postcentral Gyrus (BA 1) | 58 -28 44 | 11.874 | < 0.0001 | <0.001 | | |
| | R Superior Frontal Gyrus (BA 11) | 31 54 -17 | 11.601 | < 0.0001 | <0.001 | | |
| | L Fusiform Gyrus (BA 20) | -55 -16 -31 | 11.450 | < 0.0001 | <0.001 | | |
| | R Superior Frontal Gyrus (BA 11) | 33 60 -4 | 11.226 | < 0.0001 | <0.001 | | |
| | R Parahippocampal Gyrus (BA 37) | 31 -40 -13 | 11.002 | < 0.0001 | <0.001 | | |
| | R Inferior Temporal Gyrus (BA 20) | 52 -28 -25 | 10.657 | < 0.0001 | <0.001 | | |
| | L Lentiform Nucleus (Putamen) | -31 -20 -2 | 10.653 | < 0.0001 | <0.001 | | |
| | R Fusiform Gyrus (BA 20) | 43 -17 -31 | 10.632 | < 0.0001 | <0.001 | | |
| | R Anterior Cingulate (BA 24) | 1 -2 29 | 10.575 | < 0.0001 | <0.001 | | |
| | L Hippocampus (BA 20) | -28 -8 -13 | 10.545 | < 0.0001 | <0.001 | | |
| | L Fusiform Gyrus (BA 20) | -45 -19 -28 | 10.304 | < 0.0001 | <0.001 | | |
| | L Inferior Temporal Gyrus (BA 20) | -57 -4 -34 | 10.150 | < 0.0001 | <0.001 | | |
| | R Thalamus (Medial Dorsal Nucleus) | 6 -20 2 | 10.077 | < 0.0001 | <0.001 | | |
| | R Parahippocampal Gyrus (BA 20) | 30 -25 -17 | 10.010 | < 0.0001 | <0.001 | | |
| | R Thalamus | 7 -22 0 | 9.968 | < 0.0001 | <0.001 | | |
| | R Parahippocampal Gyrus (BA 20) | 31 -28 -16 | 9.964 | < 0.0001 | <0.001 | | |
| | R Fusiform Gyrus (BA 20) | 40 -28 -23 | 9.852 | < 0.0001 | <0.001 | | |
| | L Parahippocampal Gyrus (BA 36) | -30 -37 -13 | 9.798 | < 0.0001 | <0.001 | | |
| | L Fusiform Gyrus (BA 20) | -45 -31 -25 | 9.752 | < 0.0001 | <0.001 | | |
| | L Inferior Temporal Gyrus (BA 20) | -60 -34 -25 | 9.746 | < 0.0001 | <0.001 | | |
| | R Inferior Temporal Gyrus (BA 20) | 54 0 -38 | 9.536 | < 0.0001 | <0.001 | | |
| | L Cingulate Gyrus (BA 24) | -10 -10 41 | 9.260 | < 0.0001 | <0.001 | | |
| | L Middle Frontal Gyrus, Orbital Part (BA 10) | -37 57 -14 | 8.976 | < 0.0001 | <0.001 | | |
| | R Cingulate Gyrus (BA 24) | 9 -10 41 | 8.533 | < 0.0001 | <0.001 | | |
| | L Lingual Gyrus (BA 27) | -18 -44 -4 | 8.450 | < 0.0001 | <0.001 | | |
| | L Superior Frontal Gyrus (BA 11) | -30 50 -19 | 8.185 | < 0.0001 | <0.001 | | |
| | L Superior Temporal Gyrus (BA 38) | -40 14 -26 | 8.122 | < 0.0001 | <0.001 | | |
| | R Insula (BA 13) | 36 -14 18 | 8.057 | < 0.0001 | <0.001 | | |
| | L Supramarginal Gyrus (BA 40) | -64 -34 39 | 7.017 | < 0.0001 | <0.001 | | |
| | L Brainstem (Midbrain) | 0 -20 -19 | 6.736 | < 0.0001 | <0.001 | | |
| | L Culmen (BA 30) | -6 -43 -10 | 5.561 | < 0.0001 | 0.001 | | |
| | R Middle Temporal Gyrus (BA 20) | 36 2 -50 | 4.953 | < 0.0001 | 0.010 | | |
| 2 | L Lingual Gyrus (BA 18) | -19 -89 -7 | 10.965 | < 0.0001 | <0.001 | 1370 | 0.018 |
| 3 | L Declive | -15 -74 -28 | 7.152 | < 0.0001 | <0.001 | 1764 | 0.011 |
| | L Uvula | -27 -65 -31 | 7.116 | < 0.0001 | <0.001 | | |
| 4 | R Nodule | 9 -47 -37 | 5.516 | < 0.0001 | 0.001 | 253 | 0.214 |

Peaks with significance $p = 0.001$ or less, uncorrected. Peaks at minimum distance 10 mm. Clu#: cluster number; Coord. (mm.): coordinates in Montreal Neurological Institute; *p* (uncorr.): significance value, uncorrected; *p* (corr.): significance value, peak-level corrected; *k*: cluster extent (in 1.5 mm voxels); *p* (cl.): significance value, cluster-level corrected.

**Table S2. Foraging trend. Effect of time on task, negative direction.**

| Clu # | Brain area | Coord. (mm.) | *t* | *p* (uncorr.) | *p* (corr.) | *k* | *p* (cl.) |
|---|---|---|---|---|---|---|---|
| 1 | L Precentral Gyrus (BA 6) | -40 -14 56 | -31.364 | < 0.0001 | <0.001 | 96263 | <0.001 |
| | L Postcentral Gyrus (BA 1) | -52 -19 48 | -26.835 | < 0.0001 | <0.001 | | |
| | R Culmen (BA 37) | 22 -52 -22 | -23.574 | < 0.0001 | <0.001 | | |
| | L Rolandic Operculum | -48 -22 20 | -21.693 | < 0.0001 | <0.001 | | |
| | R Precentral Gyrus (BA 6) | 40 -13 53 | -20.990 | < 0.0001 | <0.001 | | |
| | L Medial Frontal Gyrus (BA 6) | -6 -2 57 | -19.793 | < 0.0001 | <0.001 | | |
| | L Insula (BA 6) | -42 -2 15 | -18.840 | < 0.0001 | <0.001 | | |
| | L Middle Temporal Gyrus (BA 19) | -43 -74 6 | -18.606 | < 0.0001 | <0.001 | | |
| | L Inferior Frontal Gyrus (BA 9) | -60 6 32 | -15.103 | < 0.0001 | <0.001 | | |
| | L Middle Occipital Gyrus (BA 19) | -34 -74 -14 | -14.232 | < 0.0001 | <0.001 | | |
| | R Precentral Gyrus (BA 6) | 36 -23 68 | -14.128 | < 0.0001 | <0.001 | | |
| | L Superior Parietal Lobule (BA 7) | -27 -52 68 | -13.648 | < 0.0001 | <0.001 | | |
| | R Culmen (BA 18) | 7 -59 -14 | -12.130 | < 0.0001 | <0.001 | | |
| | L Precuneus | -6 -58 72 | -12.099 | < 0.0001 | <0.001 | | |
| | R Insula | 40 -1 17 | -11.940 | < 0.0001 | <0.001 | | |
| | L Precuneus | -3 -52 74 | -11.409 | < 0.0001 | <0.001 | | |
| | L Culmen (BA 37) | -25 -53 -22 | -11.248 | < 0.0001 | <0.001 | | |
| | R Lingual Gyrus | 7 -94 -19 | -10.950 | < 0.0001 | <0.001 | | |
| | R Cuneus (BA 17) | 1 -100 -5 | -10.803 | < 0.0001 | <0.001 | | |
| | R Tuber | 52 -52 -34 | -10.668 | < 0.0001 | <0.001 | | |
| | L Declive | -51 -59 -28 | -10.603 | < 0.0001 | <0.001 | | |
| | L Calcarine (BA 17) | -1 -103 0 | -10.309 | < 0.0001 | <0.001 | | |
| | R Cerebellar Crus I | 36 -85 -25 | -10.192 | < 0.0001 | <0.001 | | |
| | L Precuneus | -3 -47 75 | -10.124 | < 0.0001 | <0.001 | | |
| | R Fusiform Gyrus (BA 18) | 24 -91 -23 | -10.063 | < 0.0001 | <0.001 | | |
| | R Declive | 48 -59 -29 | -9.845 | < 0.0001 | <0.001 | | |
| | L Paracentral Lobule | -1 -40 78 | -9.755 | < 0.0001 | <0.001 | | |
| | L Uvula (BA 18) | -12 -92 -23 | -9.668 | < 0.0001 | <0.001 | | |
| | R Precuneus (BA 5) | 1 -44 75 | -9.417 | < 0.0001 | <0.001 | | |
| | L Precuneus (BA 5) | -7 -50 78 | -9.366 | < 0.0001 | <0.001 | | |
| | R Pyramis | 3 -65 -34 | -9.332 | < 0.0001 | <0.001 | | |
| | L Cuneus (BA 19) | -21 -91 18 | -9.311 | < 0.0001 | <0.001 | | |
| | L Caudate | -18 5 26 | -8.884 | < 0.0001 | <0.001 | | |

| | | | | | | | |
|---|---|---|---|---|---|---|---|
| | L Paracentral Lobule | -3 -32 80 | -8.640 | < 0.0001 | <0.001 | | |
| | R Caudate | 19 14 21 | -8.559 | < 0.0001 | <0.001 | | |
| | R Superior Parietal Lobule (BA 7) | 16 -67 68 | -8.199 | < 0.0001 | <0.001 | | |
| | R Cuneus (BA 18) | 6 -98 12 | -8.167 | < 0.0001 | <0.001 | | |
| | R Superior Parietal Lobule (BA 7) | 19 -58 62 | -7.294 | < 0.0001 | <0.001 | | |
| | R Paracentral Lobule (BA 5) | 4 -43 78 | -6.981 | < 0.0001 | <0.001 | | |
| | R Supramarginal Gyrus (BA 1) | 27 -38 45 | -6.663 | < 0.0001 | <0.001 | | |
| | R Postcentral Gyrus | 54 -16 20 | -6.550 | < 0.0001 | <0.001 | | |
| | R Cingulate Gyrus | 19 -1 29 | -6.544 | < 0.0001 | <0.001 | | |
| | R Primary Sensory (BA 1) | 25 -40 44 | -6.506 | < 0.0001 | <0.001 | | |
| | L Precuneus | -1 -79 51 | -6.487 | < 0.0001 | <0.001 | | |
| | R Inferior Frontal Gyrus (BA 9) | 63 9 29 | -6.440 | < 0.0001 | <0.001 | | |
| | L Cingulate Gyrus (BA 6) | -12 -22 45 | -6.424 | < 0.0001 | <0.001 | | |
| | L Paracentral Lobule | -1 -22 80 | -6.272 | < 0.0001 | <0.001 | | |
| | R Precuneus (BA 5) | 6 -46 78 | -5.940 | < 0.0001 | <0.001 | | |
| | R Insula | 45 -20 21 | -5.475 | < 0.0001 | 0.001 | | |
| | R Hippocampus (BA 20) | 39 -19 -16 | -5.472 | < 0.0001 | 0.001 | | |
| | R Rolandic Operculum | 42 -22 23 | -5.422 | < 0.0001 | 0.001 | | |
| | L Precuneus (BA 7) | -6 -79 54 | -4.499 | < 0.0001 | 0.067 | | |
| | L Insula (BA 13) | -37 -5 0 | -4.049 | < 0.0001 | 0.270 | | |
| 2 | L Lentiform Nucleus (Medial Globus Pallidus) | -18 -7 -5 | -11.080 | < 0.0001 | <0.001 | 4582 | 0.003 |
| | R Hypothalamus (BA 25) | 6 0 -14 | -9.791 | < 0.0001 | <0.001 | | |
| | R Middle Temporal Pole (BA 36) | 22 12 -34 | -7.914 | < 0.0001 | <0.001 | | |
| | L Superior Temporal Gyrus (BA 38) | -24 12 -37 | -6.947 | < 0.0001 | <0.001 | | |
| | R Lentiform Nucleus (Medial Globus Pallidus) | 18 -5 -5 | -6.887 | < 0.0001 | <0.001 | | |
| | L Superior Temporal Gyrus (BA 38) | -28 18 -34 | -5.899 | < 0.0001 | <0.001 | | |
| | L Orbital Gyrus (BA 11) | -16 36 -26 | -5.309 | < 0.0001 | 0.002 | | |
| | L Superior Temporal Gyrus (BA 38) | -31 20 -32 | -4.945 | < 0.0001 | 0.012 | | |
| | R Superior Temporal Gyrus (BA 38) | 28 18 -32 | -3.903 | < 0.0001 | 0.391 | | |
| | L Inferior Frontal Gyrus (BA 11) | -22 29 -25 | -3.869 | < 0.0001 | 0.427 | | |
| | R Superior Temporal Gyrus (BA 38) | 31 20 -34 | -3.750 | 0.0001 | 0.538 | | |
| 3 | L Rectus (BA 11) | -3 66 -16 | -8.721 | < 0.0001 | <0.001 | 940 | 0.033 |
| | R Superior Frontal Gyrus, Orbital Part (BA 11) | 10 68 -16 | -6.609 | < 0.0001 | <0.001 | | |
| 4 | R Orbital Gyrus (BA 11) | 16 35 -28 | -6.924 | < 0.0001 | <0.001 | 90 | 0.526 |
| 5 | L Middle Frontal Gyrus (BA 9) | -40 48 32 | -5.267 | < 0.0001 | 0.003 | 215 | 0.255 |
| | L Middle Frontal Gyrus (BA 45) | -46 48 21 | -3.398 | 0.0004 | 0.856 | | |
| 6 | L Superior Temporal Gyrus (BA 38) | -52 18 -22 | -5.161 | < 0.0001 | 0.004 | 19 | 0.870 |
| 7 | L Substantia Nigra | -10 -20 -11 | -5.111 | < 0.0001 | 0.006 | 197 | 0.281 |

Peaks with significance $p$ = 0.001 or less, uncorrected. Peaks at minimum distance 10 mm. Clu#: cluster number; Coord. (mm.): coordinates in Montreal Neurological Institute; $p$ (uncorr.): significance value, uncorrected; $p$ (corr.): significance value, peak-level corrected; $k$: cluster extent (in 1.5 mm voxels); $p$ (cl.): significance value, cluster-level corrected.

**Table S3. Foraging trend high vs low. Interaction between time on task and reward, positive direction.**

| Clu # | Brain area | Coord. (mm.) | *t* | *p* (uncorr.) | *p* (corr.) | *k* | *p* (cl.) |
|---|---|---|---|---|---|---|---|
| 1 | R Medial Frontal Gyrus (BA 32) | 4 41 23 | 14.041 | < 0.0001 | <0.001 | 155061 | <0.001 |
| | R Insula (BA 47) | 30 21 -8 | 12.780 | < 0.0001 | <0.001 | | |
| | R Inferior Frontal Gyrus (BA 38) | 45 21 -13 | 12.690 | < 0.0001 | <0.001 | | |
| | R Anterior Cingulate (BA 32) | 7 42 5 | 12.191 | < 0.0001 | <0.001 | | |
| | R Inferior Frontal Gyrus (BA 47) | 28 18 -13 | 12.175 | < 0.0001 | <0.001 | | |
| | R Lingual Gyrus (BA 18) | 16 -71 -7 | 11.612 | < 0.0001 | <0.001 | | |
| | R Middle Temporal Gyrus (BA 21) | 58 -40 -4 | 11.270 | < 0.0001 | <0.001 | | |
| | R Middle Temporal Gyrus (BA 20) | 61 -26 -13 | 10.961 | < 0.0001 | <0.001 | | |
| | L Insula (BA 47) | -30 18 -16 | 10.839 | < 0.0001 | <0.001 | | |
| | R Medial Frontal Gyrus (BA 9) | 9 41 42 | 10.768 | < 0.0001 | <0.001 | | |
| | R Parahippocampal Gyrus (Amygdala) | 21 -4 -13 | 10.736 | < 0.0001 | <0.001 | | |
| | R Anterior Cingulate (BA 24) | 4 27 18 | 10.576 | < 0.0001 | <0.001 | | |
| | R Inferior Parietal Lobule (BA 40) | 49 -49 47 | 10.573 | < 0.0001 | <0.001 | | |
| | R Superior Frontal Gyrus (BA 6) | 7 29 59 | 10.555 | < 0.0001 | <0.001 | | |
| | R Superior Frontal Gyrus (BA 10) | 16 56 24 | 10.465 | < 0.0001 | <0.001 | | |
| | R Middle Temporal Gyrus (BA 21) | 55 -28 -5 | 10.463 | < 0.0001 | <0.001 | | |
| | R Inferior Frontal Gyrus (BA 45) | 52 23 23 | 10.462 | < 0.0001 | <0.001 | | |
| | R Superior Frontal Gyrus (BA 9) | 13 48 36 | 10.298 | < 0.0001 | <0.001 | | |
| | L Declive | -27 -68 -28 | 9.669 | < 0.0001 | <0.001 | | |
| | R Middle Frontal Gyrus (BA 10) | 24 62 9 | 9.272 | < 0.0001 | <0.001 | | |
| | R Precuneus (BA 19) | 16 -83 38 | 9.144 | < 0.0001 | <0.001 | | |
| | R Middle Frontal Gyrus (BA 9) | 43 12 50 | 9.011 | < 0.0001 | <0.001 | | |
| | L Declive | -10 -80 -28 | 8.874 | < 0.0001 | <0.001 | | |
| | R Middle Frontal Gyrus (BA 47) | 33 56 -1 | 8.804 | < 0.0001 | <0.001 | | |
| | R Middle Frontal Gyrus (BA 11) | 45 44 -13 | 8.800 | < 0.0001 | <0.001 | | |
| | R Middle Frontal Gyrus (BA 47) | 37 53 -5 | 8.688 | < 0.0001 | <0.001 | | |
| | R Cingulate Gyrus (BA 23) | 1 -16 38 | 8.517 | < 0.0001 | <0.001 | | |
| | R Precentral Gyrus (BA 9) | 46 20 39 | 8.423 | < 0.0001 | <0.001 | | |
| | R Posterior Cingulate (BA 30) | 6 -62 5 | 8.286 | < 0.0001 | <0.001 | | |
| | R Caudate (Caudate Body) | 13 8 14 | 8.238 | < 0.0001 | <0.001 | | |
| | R Hippocampus | 21 -17 -11 | 8.187 | < 0.0001 | <0.001 | | |
| | R Thalamus | 7 -5 8 | 7.992 | < 0.0001 | <0.001 | | |

| | | | | |
|---|---|---|---|---|
| R Precuneus (BA 7) | 10 -79 47 | 7.858 | < 0.0001 | <0.001 |
| R Cingulate Gyrus (BA 23) | 7 -40 38 | 7.820 | < 0.0001 | <0.001 |
| R Middle Frontal Gyrus (BA 46) | 42 39 21 | 7.404 | < 0.0001 | <0.001 |
| R Cuneus (BA 18) | 6 -79 35 | 7.214 | < 0.0001 | <0.001 |
| R Middle Temporal Gyrus (BA 37) | 52 -58 -14 | 7.010 | < 0.0001 | <0.001 |
| R Parahippocampal Gyrus (BA 27) | 16 -40 -4 | 6.926 | < 0.0001 | <0.001 |
| R Cuneus (BA 18) | 7 -86 23 | 6.898 | < 0.0001 | <0.001 |
| R Middle Temporal Gyrus (BA 37) | 54 -53 -13 | 6.839 | < 0.0001 | <0.001 |
| R Declive (BA 19) | 31 -70 -26 | 6.809 | < 0.0001 | <0.001 |
| R Parahippocampal Gyrus (BA 19) | 19 -44 -7 | 6.707 | < 0.0001 | <0.001 |
| L Superior Frontal Gyrus (BA 10) | -25 51 -2 | 6.582 | < 0.0001 | <0.001 |
| L Inferior Frontal Gyrus (BA 47) | -51 36 -13 | 6.572 | < 0.0001 | <0.001 |
| R Calcarine (BA 19) | 22 -55 8 | 6.546 | < 0.0001 | <0.001 |
| L Anterior Cingulate (BA 24) | -3 23 -5 | 6.530 | < 0.0001 | <0.001 |
| L Inferior Frontal Gyrus (BA 45) | -58 20 12 | 6.529 | < 0.0001 | <0.001 |
| L Lingual Gyrus (BA 18) | -6 -62 2 | 6.500 | < 0.0001 | <0.001 |
| L Caudate (Caudate Head) | -9 6 0 | 6.469 | < 0.0001 | <0.001 |
| L Tuber | -37 -76 -34 | 6.389 | < 0.0001 | <0.001 |
| L Culmen (BA 37) | -18 -49 -19 | 6.334 | < 0.0001 | <0.001 |
| R Lentiform Nucleus (Putamen) | 18 9 -1 | 6.286 | < 0.0001 | <0.001 |
| L Superior Frontal Gyrus (BA 8) | -10 27 56 | 6.232 | < 0.0001 | <0.001 |
| L Culmen (BA 18) | -13 -55 -13 | 6.170 | < 0.0001 | <0.001 |
| L Parahippocampal Gyrus (BA 34) | -18 -2 -13 | 6.121 | < 0.0001 | <0.001 |
| R Inferior Temporal Gyrus (BA 37) | 43 -49 -10 | 5.909 | < 0.0001 | <0.001 |
| L Middle Frontal Gyrus (BA 47) | -34 54 -7 | 5.896 | < 0.0001 | <0.001 |
| L Superior Frontal Gyrus (BA 9) | -19 44 32 | 5.841 | < 0.0001 | <0.001 |
| R Superior Frontal Gyrus (BA 11) | 16 54 -10 | 5.708 | < 0.0001 | 0.001 |
| L Posterior Cingulate (BA 17) | -21 -59 9 | 5.560 | < 0.0001 | 0.001 |
| L Cerebellar Tonsil | -7 -55 -41 | 5.530 | < 0.0001 | 0.001 |
| L Superior Frontal Gyrus (BA 46) | -19 53 21 | 5.489 | < 0.0001 | 0.002 |
| R Brainstem (Midbrain) | 7 -14 -11 | 5.386 | < 0.0001 | 0.002 |
| R Declive (BA 18) | 12 -79 -23 | 5.304 | < 0.0001 | 0.004 |
| R Inferior Frontal Gyrus (BA 45) | 36 29 11 | 5.139 | < 0.0001 | 0.006 |
| L Inferior Frontal Gyrus (BA 45) | -43 21 6 | 5.004 | < 0.0001 | 0.012 |
| L Uvula | -6 -70 -37 | 4.932 | < 0.0001 | 0.016 |
| R Parahippocampal Gyrus (BA 37) | 28 -37 -13 | 4.755 | < 0.0001 | 0.031 |
| R Precuneus (BA 5) | 3 -56 68 | 4.705 | < 0.0001 | 0.036 |
| L Posterior Cingulate (BA 30) | -15 -67 11 | 4.676 | < 0.0001 | 0.041 |
| L Lingual Gyrus (BA 19) | -15 -47 -5 | 4.607 | < 0.0001 | 0.053 |
| R Middle Temporal Gyrus (BA 20) | 54 -7 -16 | 4.182 | < 0.0001 | 0.228 |

| | | | | | | | |
|---|---|---|---|---|---|---|---|
| | L Cerebellar Tonsil | -36 -61 -44 | 4.155 | < 0.0001 | 0.247 | | |
| | L Pyramis | -52 -65 -43 | 4.070 | < 0.0001 | 0.317 | | |
| | R Thalamus | 6 -26 0 | 3.760 | < 0.0001 | 0.615 | | |
| | L Precentral Gyrus (BA 9) | -39 21 41 | 3.727 | 0.0001 | 0.650 | | |
| | L Cerebellar Tonsil | -19 -43 -47 | 3.670 | 0.0001 | 0.708 | | |
| | L Middle Frontal Gyrus (BA 9) | -39 9 47 | 3.585 | 0.0002 | 0.785 | | |
| | L Culmen | -28 -35 -35 | 3.402 | 0.0004 | 0.921 | | |
| 2 | L Middle Temporal Gyrus (BA 21) | -67 -25 -10 | 6.787 | < 0.0001 | <0.001 | 2743 | 0.004 |
| | L Middle Temporal Gyrus (BA 21) | -61 -32 -7 | 6.308 | < 0.0001 | <0.001 | | |
| | L Inferior Temporal Gyrus (BA 20) | -48 -52 -13 | 4.492 | < 0.0001 | 0.082 | | |
| 3 | L Supramarginal Gyrus (BA 22) | -58 -52 24 | 6.302 | < 0.0001 | <0.001 | 4499 | 0.001 |
| | L Supramarginal Gyrus (BA 39) | -49 -53 32 | 5.786 | < 0.0001 | <0.001 | | |
| | L Inferior Parietal Lobule (BA 40) | -48 -55 47 | 5.554 | < 0.0001 | 0.001 | | |
| | L Inferior Parietal Lobule (BA 40) | -66 -40 21 | 3.811 | < 0.0001 | 0.556 | | |
| 4 | L Cerebellar Crus II | -48 -46 -41 | 5.084 | < 0.0001 | 0.008 | 273 | 0.226 |

Peaks with significance *p* = 0.001 or less, uncorrected. Peaks at minimum distance 10 mm. Clu#: cluster number; Coord. (mm.): coordinates in Montreal Neurological Institute; *p* (uncorr.): significance value, uncorrected; *p* (corr.): significance value, peak-level corrected; *k*: cluster extent (in 1.5 mm voxels); *p* (cl.): significance value, cluster-level corrected.

**Table S4. Foraging trend high vs low. Interaction between time on task and reward, negative direction.**

| Clu # | Brain area | Coord. (mm.) | *t* | *p* (uncorr.) | *p* (corr.) | *k* | *p* (cl.) |
|---|---|---|---|---|---|---|---|
| 1 | L Precentral Gyrus (BA 6) | -31 -22 71 | -12.729 | < 0.0001 | <0.001 | 25158 | <0.001 |
| | L Precentral Gyrus (BA 6) | -39 -19 65 | -12.572 | < 0.0001 | <0.001 | | |
| | L Precentral Gyrus (BA 4) | -39 -16 48 | -10.418 | < 0.0001 | <0.001 | | |
| | L Postcentral Gyrus (BA 43) | -49 -14 17 | -7.668 | < 0.0001 | <0.001 | | |
| | L Superior Temporal Gyrus (BA 22) | -60 -11 2 | -6.328 | < 0.0001 | <0.001 | | |
| | L Precentral Gyrus (BA 6) | -52 -4 26 | -5.841 | < 0.0001 | <0.001 | | |
| | L Cingulate Gyrus (BA 24) | -9 5 48 | -5.724 | < 0.0001 | <0.001 | | |
| | L Superior Frontal Gyrus (BA 6) | -18 2 72 | -5.044 | < 0.0001 | 0.010 | | |
| | L Caudate | -19 17 20 | -4.838 | < 0.0001 | 0.023 | | |
| | R Medial Frontal Gyrus (BA 6) | 9 2 53 | -4.728 | < 0.0001 | 0.035 | | |
| | L Caudate | -18 8 26 | -4.593 | < 0.0001 | 0.055 | | |
| | L Cingulate Gyrus | -16 -22 44 | -4.576 | < 0.0001 | 0.058 | | |
| | L Medial Frontal Gyrus (BA 6) | -4 -7 59 | -4.335 | < 0.0001 | 0.138 | | |
| | L Precuneus (BA 7) | -19 -50 56 | -4.089 | < 0.0001 | 0.286 | | |
| | L Inferior Parietal Lobule (BA 5) | -28 -43 54 | -4.031 | < 0.0001 | 0.338 | | |
| | L Inferior Frontal Gyrus (BA 9) | -60 6 36 | -3.908 | < 0.0001 | 0.456 | | |
| 2 | R Culmen (BA 37) | 19 -50 -22 | -8.286 | < 0.0001 | <0.001 | 1415 | 0.014 |
| 3 | L Lingual Gyrus (BA 18) | -13 -79 -7 | -8.067 | < 0.0001 | <0.001 | 1461 | 0.013 |

| | | | | | | | |
|---|---|---|---|---|---|---|---|
| 4 | L Inferior Temporal Gyrus (BA 20) | -37 -7 -37 | -5.877 | < 0.0001 | <0.001 | 2488 | 0.004 |
| | L Fusiform Gyrus (BA 36) | -28 2 -38 | -4.986 | < 0.0001 | 0.013 | | |
| | L Fusiform Gyrus (BA 20) | -52 -37 -28 | -4.145 | < 0.0001 | 0.243 | | |
| | L Parahippocampal Gyrus (BA 20) | -31 -20 -26 | -3.533 | 0.0002 | 0.825 | | |
| 5 | R Uncus (BA 20) | 28 2 -41 | -5.653 | < 0.0001 | <0.001 | 1024 | 0.025 |
| 6 | L Cuneus (BA 19) | -19 -91 29 | -5.556 | < 0.0001 | 0.001 | 5576 | 0.001 |
| | L Middle Occipital Gyrus (BA 18) | -30 -97 6 | -5.269 | < 0.0001 | 0.003 | | |
| | L Middle Occipital Gyrus (BA 17) | -9 -103 12 | -5.254 | < 0.0001 | 0.004 | | |
| | L Primary Visual Cortex (BA 17) | -7 -106 -5 | -5.129 | < 0.0001 | 0.006 | | |
| | L Middle Occipital Gyrus (BA 19) | -46 -76 5 | -5.092 | < 0.0001 | 0.008 | | |
| | L Primary Visual Cortex (BA 17) | -1 -104 3 | -5.030 | < 0.0001 | 0.011 | | |
| | L Middle Occipital Gyrus (BA 18) | -21 -98 14 | -4.733 | < 0.0001 | 0.035 | | |
| | L Inferior Occipital Gyrus (BA 18) | -36 -92 -13 | -4.137 | < 0.0001 | 0.249 | | |
| | L Cuneus (BA 19) | -7 -91 38 | -3.770 | < 0.0001 | 0.595 | | |
| 7 | L Superior Frontal Gyrus (BA 9) | -34 47 38 | -5.135 | < 0.0001 | 0.006 | 573 | 0.076 |
| | L Middle Frontal Gyrus (BA 8) | -34 39 45 | -4.973 | < 0.0001 | 0.013 | | |
| 8 | R Precentral Gyrus (BA 43) | 61 -1 23 | -4.879 | < 0.0001 | 0.020 | 816 | 0.040 |
| | R Precentral Gyrus (BA 6) | 43 -10 38 | -4.312 | < 0.0001 | 0.148 | | |
| 9 | R Middle Occipital Gyrus (BA 18) | 34 -94 12 | -4.639 | < 0.0001 | 0.047 | 485 | 0.100 |
| | R Middle Occipital Gyrus (BA 18) | 27 -101 0 | -4.141 | < 0.0001 | 0.246 | | |
| | R Cuneus (BA 18) | 19 -104 -7 | -3.823 | < 0.0001 | 0.541 | | |

Peaks with significance $p$ = 0.001 or less, uncorrected. Peaks at minimum distance 10 mm. Clu#: cluster number; Coord. (mm.): coordinates in Montreal Neurological Institute; $p$ (uncorr.): significance value, uncorrected; $p$ (corr.): significance value, peak-level corrected; $k$: cluster extent (in 1.5 mm voxels); $p$ (cl.): significance value, cluster-level corrected.
